# Ancient genomic insights into *Salmonella enterica* Paratyphi C from Central Mexico

**DOI:** 10.1101/2025.10.13.681940

**Authors:** Miriam Bravo-Lopez, Eduardo Arrieta-Donato, Viridiana Villa-Islas, Åshild Joanne Vågene, Ana B. Villaseñor-Altamirano, Ernesto Garfias Morales, Laura Carrillo-Olivas, Jorge Gómez-Valdés, Sandra Elena Guevara Flores, Alejandro Meraz Moreno, Maria Moreno Cabrera, Jean Cury, Flora Jay, Emilia Huerta-Sanchez, María C. Ávila-Arcos

## Abstract

*Salmonella enterica* is a widespread pathogen of major global health relevance, with over 2,500 serovars classified into non-typhoidal and typhoidal groups. Within the typhoidal group, *S. enterica* Paratyphi C causes paratyphoid fever in humans. Ancient DNA from this bacterium has previously been recovered from epidemic-associated burials in Eurasia and Mexico, dating back 6,000 to 300 years. Here, we analyzed dental DNA from seven individuals (radiocarbon dated to 1800–1940 CE) buried at the Temple of the Immaculate Conception in Mexico City, Central Mexico, and identified ancient *S. enterica* Paratyphi C DNA in a young female with Native American ancestry. Using an in-house targeted enrichment strategy and deep shotgun sequencing, we reconstructed a *S. enterica* Paratyphi C genome (COYC5) with ∼11X coverage. Phylogenetic and comparative genomic analyses place COYC5 in close association with previously reported *S. enterica* Paratyphi C genomes associated with the 1545 *cocoliztli* epidemic in southern Mexico, as well as with European strains. Divergence estimates indicate that the Mexican and European lineages shared a common ancestor approximately 1,400 years before present (yBP), reflecting an ancient evolutionary split predating European colonization of Mexico. In contrast, the divergence between COYC5 and the southern Mexican genomes occurred around 516 yBP, coinciding with the onset of the colonial period. This pattern supports a European introduction of *S. enterica* Paratyphi C during colonization, followed by its local diversification within Mexico. Despite this regional differentiation, the conserved presence of key virulence loci—such as SPI-7 and an active shufflon system—across COYC5, southern Mexican, and European genomes underscores the enduring pathogenic potential of *S. enterica* Paratyphi C. The identification of this bacterium in 19th-century Mexico City provides the first genomic evidence of its persistence in urban contexts beyond major epidemic outbreaks, offering new insights into its evolutionary trajectory in Mexico. These findings raise new questions about how the pathogen spread and persisted across different ecological, social, and epidemiological contexts in ancient Mexico and the Americas.

## 1. Introduction

*Salmonella enterica* is classified by the World Health Organization (WHO) as a high-priority bacterial pathogen due to its global public health impact (WHO, 2024). This species comprises more than 2,579 serovars, which are divided into typhoidal *Salmonella* (TS)—including serovars Typhi and Paratyphi A, B, and C—and non-typhoidal *Salmonella* (NTS), such as *S.* Enteritidis, *S.* Typhimurium, *S.* Heidelberg, and *S.* Newport. In 2019, the WHO estimated that NTS caused over 500,000 cases of invasive disease and more than 79,000 deaths worldwide, while TS accounted for approximately 9 million cases and 110,000 deaths (WHO, 2023). Overall, infections with *S. enterica* are thought to result in between 100 million cases and 200,000 deaths annually, underscoring its importance as a global health threat (Lamichhane et al., 2024).

Although NTS and TS share over 97% of their genes, they exhibit distinct disease manifestations (Liu et al., 2009; Sabbagh et al., 2010). NTS typically causes gastroenteritis of milder severity than that associated with TS and is most transmitted through contaminated food. However, in immunocompromised individuals, NTS infections may progress to severe disease (Kumar et al., 2025).

In contrast, TS serovars exclusively infect humans and, if left untreated, can have a mortality rate of up to 30% (Wang et al., 2023). While typhoid fever has declined in most developed countries, it remains a significant public health concern in many low- and middle-income countries (Als et al., 2018). TS disseminates systemically through mesenteric lymph nodes to the liver and spleen, resulting in symptoms such as fever, diarrhea, nausea, and abdominal pain, depending on the host’s immune status (Gal-Mor et al., 2014; Hiyoshi et al., 2018).

Among the typhoidal serovars, *S. enterica* Paratyphi A is the most frequently reported in modern human populations, followed by Paratyphi B and Paratyphi C (Als et al., 2018). Although *S. enterica* Paratyphi C causes a typhoid-like disease in humans, it is proposed to have diverged from a common ancestor with *S. Choleraesuis* through the accumulation of genomic adaptations during its evolution in human hosts (Liu et al., 2009). Overall, *Salmonella* has demonstrated remarkable adaptability to diverse host niches and environments, driven not only by the acquisition of genes via horizontal transfer but also by the loss of genes or gene functions (Han et al., 2024)

Paleogenomic evidence places *S. enterica* Paratyphi C in human populations across Eurasia and Mexico over a span of nearly six millennia (∼6,000 to 300 years ago), recovered from diverse ecological settings and archaeological contexts. The earliest evidence of this pathogen, dating to ∼6,000 years BP in Eurasia, places it within the Ancient Eurasian Super Branch (AESB), a diverse group of closely related serovars, which comprises the human-specific serovar Paratyphi C and a broad range of animal-adapted *S. enterica* serovars (Key et al., 2020). Host adaptation within the AESB likely coincided with the Neolithic transition, whose ecological context facilitated sustained exposure to *S. enterica* (Key et al., 2020). The Proto-Silk Road further contributed to the spread of *S. enterica* Paratyphi C, with evidence of its presence in Eastern Eurasia dating back at least 3,000 years. The marked differences of two Bronze Age genomes from China suggest genetic heterogeneity, with strains associated with systemic paratyphoid-like infections and non-typhoidal salmonellosis (Wu et al., 2021). By 2,000 yBP, *S. enterica* Paratyphi C had expanded into the eastern Mediterranean, where host-generalist strains emerged, capable of infecting diverse species (Neumann et al., 2022). Its persistence across millennia highlights its adaptability to different hosts and cultural settings.

The epidemic potential of *S. enterica* Paratyphi C is documented in the medieval period, with outbreaks in 14th-century Norway and Germany (Haller, Callan, et al., 2021; Zhou et al., 2018). Ancient DNA (aDNA) from this bacterium was also identified in 16th-century individuals from Mexico associated with the *cocoliztli* epidemic (1545–1550) in Teposcolula-Yucundaa, southern Mexico (Vågene et al., 2018). However, its occurrence in other regions and in non-epidemic contexts in Mexico has remained largely unexplored.

Within this context, the identification of aDNA from *S. enterica* Paratyphi C in individuals from a 19th-century site in Mexico City—namely, The Temple of the Immaculate Conception (“La Conchita”), where the archaeological context is not linked to epidemic events—prompted us to explore in depth its phylogenetic relationship with other ancient and modern strains across its global distribution. To this end, we reconstructed a ∼11X coverage genome of *S. enterica* Paratyphi C (COYC5) using an in-house hybridization capture-enrichment protocol and deep sequencing. The reconstructed genome allowed us to define a distinct ancient Mexican sublineage including COYC5 and the previously reported ancient *S. enterica* genomes from southern Mexico. The phylogenetic position and divergence dating estimates suggest an introduction to Mexico during European colonization, following local diversification.

## 2. Results

We reanalyzed the paleogenomic data from teeth samples of seven individuals excavated at the Temple of the Immaculate Conception in Mexico City, dating from 1800 to 1899 CE, previously reported in Bravo-Lopez *et al.,* (2020) and Guzmán-Solís *et al.,* (2021) (Table S1). The paleogenomic data from the seven individuals consisted of approximately 2 million to 245 million paired-end reads per sample.

### 2.1. Identification of *Salmonella enterica* ancient DNA in one individual from the Temple of the Immaculate Conception

The metagenomic characterization of the unmapped reads to the human reference genome was performed using Kraken2 (Wood et al., 2019). The taxonomic profiles of the seven individuals revealed the presence of predominantly environmental and human oral bacteria at varying abundances (Table S2), confirming previous findings in these individuals of aDNA belonging to *T. forsythia* and Human Parvovirus B19 (Bravo-Lopez et al., 2020; Guzmán-Solís et al., 2021).

Interestingly, one individual, COYC5, had 11,235 reads assigned to *Salmonella enterica* according to Kraken2 (Table S2). Additionally, the reads from the DNA extraction and library construction negative controls assigned no reads to *S. enterica* (Table S2), supporting the authenticity of the result. As the Kraken2 database did not include genomes from *S. enterica* subspecies, we applied a competitive mapping strategy to determine the specific subspecies present. This analysis included both human-restricted and generalist *S. enterica* serovars (see Methods). The highest number of uniquely mapped reads (1,864) corresponded to the *S. enterica* Paratyphi C RKS4594 genome (NC_012125.1), identifying it as the most likely infecting serovar (Table S3).

After aligning the sequencing reads from the COYC5 individual against the *S. enterica* Paratyphi C RKS4594 genome, 48,039 reads mapped to the chromosome, of which 1,215 reads aligned to the virulence plasmid pSPCV (Table S4). These reads met the following aDNA authentication criteria: (i) high-frequency deamination at the 5′ and 3′ ends; (ii) homogeneous distribution across the reference genome; and (iii) a decreasing edit distance pattern (Figures S3 and S4).

Notably, the osteological assessment of the COYC5 skull—the only skeletal element recovered— did not reveal a specific pathological condition. The individual was identified as a female of approximately 14 years of age (Figure S1). The remains correspond to a secondary deposition within a simple burial pit (Figure S2) and were radiocarbon (C14) and contextually dated to 1800-1899 CE (Figure S5, See Methods). Evidence of physiological stress was minimal, limited to slight enamel hypoplasia, mild alveolar resorption, and periodontal inflammation. Additionally, subtle lipping of the occipital condyles was observed, likely reflecting axial loading stress associated with physically demanding activity.

### 2.2. Reconstruction of the S. *enterica* Paratyphi C genome through an in-house hybridization capture

To enhance the coverage of *S. enterica* Paratyphi C in the COYC5 individual, we implemented an in-house whole-genome capture-enrichment strategy involving the construction of biotinylated RNA baits (Carpenter et al., 2013). This included generating biotinylated RNA baits transcribed from the DNA of a modern *S. enterica* serovar Typhimurium strain SO2 (see Methods), a non-typhoidal serovar isolated and genomically characterized from a Mexican individual (Silva et al., 2016). While genomes of typhoidal and non-typhoidal *Salmonella enterica* serovars share more than 97% sequence similarity (Liu et al., 2009), they differ markedly in their pathogenicity islands (SPIs), which contain many of the key virulence factors, including the virulence plasmid pSPCV— which was not directly targeted by the baits.

To overcome the limitations of our probe-based capture, we complemented our enrichment approach with deep shotgun sequencing (∼245 million reads). This strategy enabled the recovery of genomic regions that were poorly captured due to potential biases in probe design. As shown in Table S5, we downsampled the same number of reads from the pre-capture, capture, and deep sequencing + capture libraries to compare the recovery of the pSPCV plasmid. The pre-capture (shotgun) library yielded 297 mapped reads, covering 29.89% of the plasmid, while the capture-only library had 46 mapped reads, covering 5.13%. In particular, the combined deep sequencing + capture library recovered 297 reads and 28.39% coverage—nearly identical to the pre-capture library. These results suggest that the combined use of capture and deep sequencing enabled the recovery of Paratyphi C key virulence factors, such as the pSPCV plasmid.

The capture-enriched library yielded a 5.2-fold increase in target sequence representation compared to pre-capture libraries (Table S5, Figure S4), with endogenous DNA content rising from 0.50% to 2.58%. As expected for capture-enrichment experiments, clonality also increased, with 90% of reads being duplicates in the enriched library, compared to 0.67% prior to capture (Table S5). Enrichment on- and off-target was subsequently assessed, and a chi-squared test indicated a highly significant difference (χ² = 34,550, df = 1, p < 2.2 × 10e-16, see Methods). Overall, this approach enabled reconstruction of an 11X coverage *S. enterica* Paratyphi C genome.

### 2.3. COYC5 forms a distinct sublineage within Post-Contact Southern Mexico genomes

To understand the phylogenetic relationships of ancient and modern *S. enterica* Paratyphi C genomes, we built a Maximum Likelihood (ML) tree based on a multi-SNP alignment of our COYC5 genome and 219 modern genomes that belong to the “Para C Lineage” (Key et al., 2020), which includes the Paratyphi C, Typhisuis, Choleraesuis serovars, plus two Birkenhead strains, as well as the rare serovar Lomita (Key et al., 2020). Additionally, 14 ancient genomes with at least 3-fold coverage were included: Tepos 14 and Tepos 35 from southern Mexico (Vågene et al., 2018); Ragna (Zhou et al., 2018); IV3002, MUR019, MUR009, SUA004, IKI003, and ETR001 (Key et al., 2020); HGH-1600 (Haller, Callan, et al., 2021); XBQM20 and XBQM90 (Wu et al., 2021); HGC004 and HGC040 (Neumann et al., 2022) (Figure 1 and Table S7).

**Figure 1.**
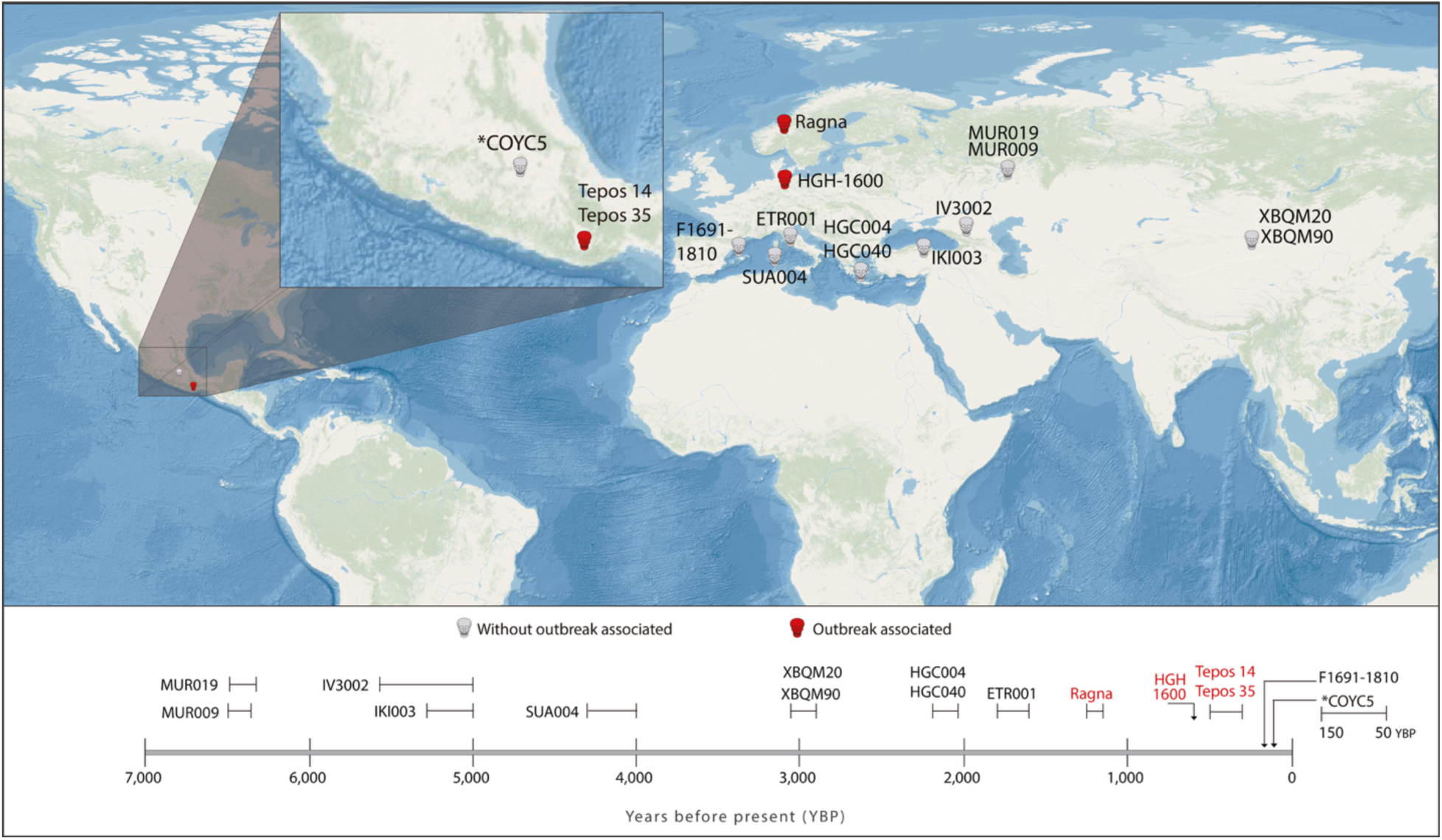
Geographic location and radiocarbon age of ancient human individuals infected with *S. enterica*. Newly sequenced genome in this study (COYC5). Previously published ancient genomes from Vågene *et al.,* 2018 (Tepos 14, and Tepos 35); Zhou *et al.,* 2018 (Ragna); and Key *et al.,* 2020 (MUR009, MUR019, IKI003, IV3002, SUA004, ETR001); de-Dios *et al*., 2021 (F1691-1810); Haller, Callan, *et al.,* 2021 (HGH1600); Neumann *et al*., 2021 (HGC004 and HGC040), and Wu *et al*., 2021 (XBQM20 and XBQM90). The red font indicates association with an epidemic outbreak.

COYC5 clusters closely with the Post-Contact southern Mexican genomes Tepos 14 and Tepos 35, although it falls within a distinct sub-branch (Figure 2). Together, these three genomes form a clade nested within the diversity of *S. enterica* Paratyphi C, specifically within the PC-1 and PC-2 clusters defined by Key et al. (2020), which comprise nearly 90 genomes from Europe and Africa (Zhou et al., 2018). In contrast, the two Pre-Contact medieval European genomes—HGH-1600 (583 yBP, Germany) and Ragna (1200 yBP, Norway)—cluster together in a basal sister clade to both the Mexican genomes and the broader set of modern *S. enterica* Paratyphi C genomes (Figure 2).

**Figure 2.**
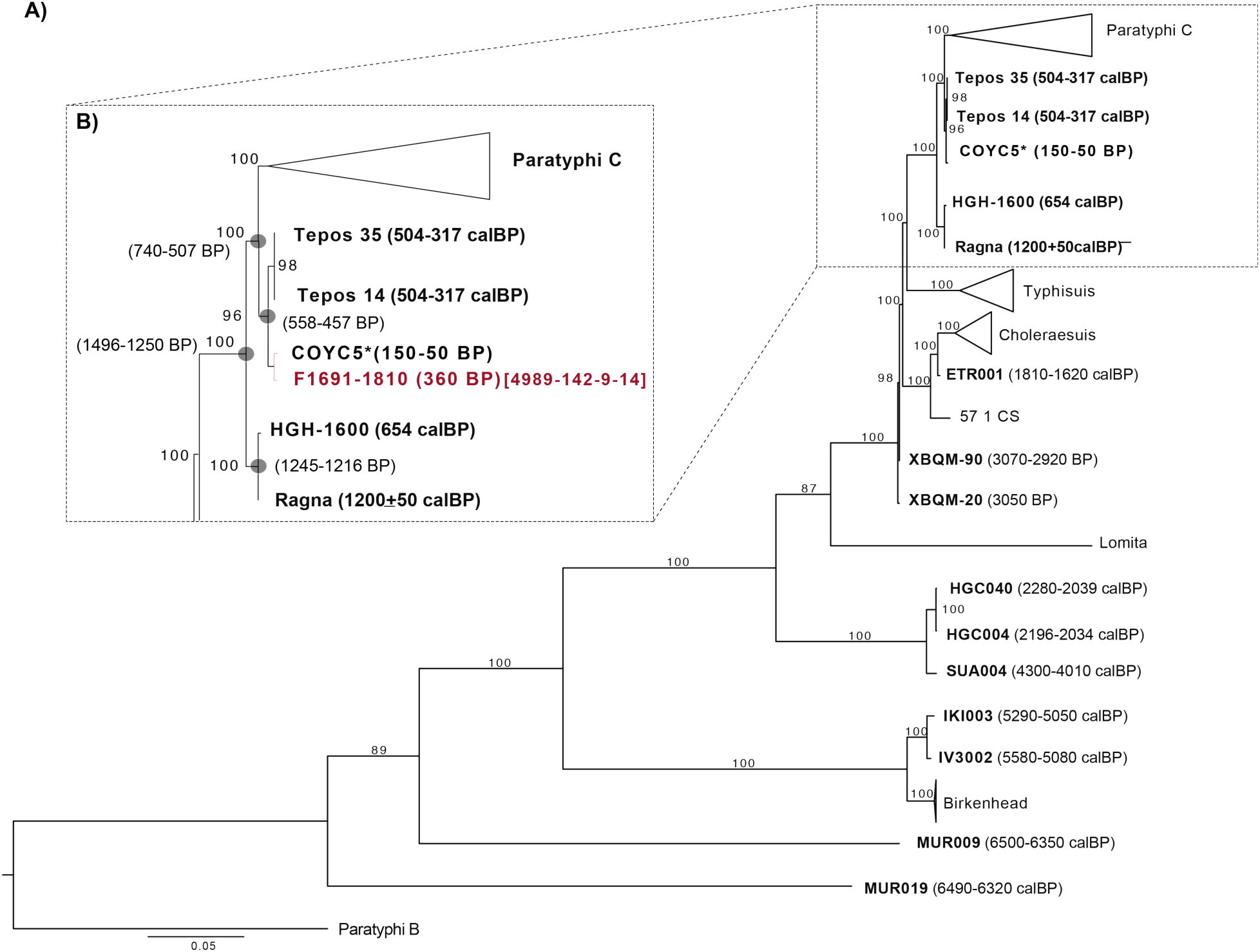
Phylogenetic relationships of reconstructed ancient and modern *S. enterica* genomes. **A.** Maximum-likelihood tree including the ancient genomes (>3X coverage), based on 93,590 SNP positions with 1,000 bootstrap replicates. Bootstrap values are shown at each node. *S. enterica* Paratyphi B SPB7 was used as the outgroup. **B**. Zoomed view of the COYC5 placement. The dates placed in the nodes indicate divergence times estimated by BEAST. Sample F1691-1810 is schematically placed based on the phylogenetic placement obtained using *pathPhynder;* the number in brackets indicates [supporting markers above the branch – conflicting markers above the branch – supporting markers on the branch – conflicting markers on the branch].

We calculated the pairwise genetic distance using the maximum-likelihood approach in RAxML (Figure S6), which reflected the clustering patterns observed in the phylogenetic tree. Notably, COYC5 shows the greatest genomic similarity to Tepos and exhibits a closer relationship to the medieval European genomes HGH-1600 (583 yBP, Germany) and Ragna (1200 BP, Norway) than to other ancient *S. enterica* Paratyphi C genomes.

### 2.4. COYC5 represents a lineage that diverged from southern Mexican genomes during the early colonial period

To further understand the temporal split between COYC5 and the genomes from southern Mexico (Tepos), we assessed the temporal signal using *TempEst* (Rambaut et al., 2016). The analysis confirmed that the SNP alignment contains a sufficient temporal signal to support a molecular clock-based phylogenetic dating (Figure S7). The regression yielded an R² value of approximately 0.32 and a p-value < 2.2e-16, demonstrating a significant correlation between genetic divergence and sampling time. Subsequently, we conducted three independent BEAST runs, a Bayesian Skyline Coalescent model, and a strict molecular clock to estimate divergence times (see Methods).

The coalescent-based BEAST analysis estimated the MRCA of the Mexican and European *S. enterica* Paratyphi C genomes at approximately 1402 BP (95% HPDI: 1496–1250 BP) (Figure 2, Figure S8). This indicates that the Mexican and European lineages share a common ancestor that diverged more than a millennium ago. Within Mexico, the divergence of the Tepos individuals and the COYC5 was estimated at 516 BP (95% HPDI: 558–457 BP). The COYC5 and Southern Mexico genomes likely descend from a common ancestor that arrived in Mexico during early colonial times, after which the lineages split and evolved separately.

The reliability of our coalescent-based estimate is supported by the concordant divergence date obtained for previously reported genomes, such as IV3002 and IKI003, which we dated to 5,656 BP (95% HPDI: 5,622.27 - 5,484.08), and is broadly consistent with the previously reported 5,681 BP (Key et al., 2020) (Figure S8).

To provide phylogenetic context and assess potential European connections, we positioned the low-coverage genome F1691-1810 (0.3X) from 17th-century Spain (de-Dios et al., 2021), within the previously obtained ML phylogeny. This individual has been shown to cluster with Post-Contact Mexican *S. enterica* Paratyphi C (de-Dios et al., 2021).

We used the program *pathPhynder* (Martiniano et al., 2022), which maps informative variants onto a reference phylogeny and assigns placement based on a four-part score (see Methods). We observed that F1691-1810 clustered adjacent to the Mexican genome COYC5, forming a distinct group (Figures 2 and S9). Its *pathPhynder* score was 4989–142–9–14, where the first two values represent supporting (4989) and conflicting (142) markers on the branch above, and the latter two indicate supporting (9) and conflicting (14) markers at the branch next to COYC5. The strong excess of support over conflict at the higher branch level confirms its assignment to the Tepos–COYC5 group, while the relatively balanced support and conflict at the lower branch level indicate uncertainty in its exact placement within this cluster.

To further evaluate the method’s reliability, we additionally placed three other low-coverage genomes (<2X): XBQ7 (3210–3000 BP, from China) and XBQ64 (undated, from China) (Wu et al. 2021), and OBP001 (5320–5070 BP, from Switzerland) (Key et al. 2020) (Figure S9). The placements of these genomes were consistent with those reported previously. The XBQ7 and XBQ64 genomes aligned closely with those previously recovered from the same archaeological site in Xinjiang, China (Wu et al., 2021). The OBP001 genome branched off basally within the Para C lineage, which includes serovars Paratyphi C, Choleraesuis, Typhisuis, and Lomita, as reported by Key et al. (2020). Altogether, these results reinforce previously published findings (Key et al., 2020; Wu et al., 2021) and support the phylogenetic placement of F1691-1810 by *pathPhynder*.

### 2.5. Host ancestry of COYC5 suggests Native American origin

The autosomal and mitochondrial genetic ancestry of the COYC5 individual was analyzed using the screening and high-depth sequencing data. In total, 10,433,726 reads were assigned to the human genome (0.28X depth of coverage), while 68,266 (414.8X depth of coverage) were assigned to the mitochondrial genome (Table S8). The molecular sex was assigned to the individual as female using the method described in Skoglund, Storå, Götherström, & Jakobsson (2013).

To explore the continental genetic ancestry of the COYC5 individual, we used the 1000 Genomes Project (1KGP) (Consortium & The 1000 Genomes Project Consortium, 2015) as a reference panel. First, we selected 10 populations as representatives of the continental superpopulations, with African (YRI, ESN, GWD, MSL), European (IBS, CEU), East Asian (CHB, CHS), and Native American (MXL, PEL) ancestry.

A Principal Components Analysis (PCA) shows that COYC5 clusters with present-day Mexican and Peruvian populations, specifically with those closest to the East Asian populations, which is consistent with a higher proportion of Native American genetic ancestry (Fig. 3A). The ADMIXTURE analysis indicates that COYC5’s host has solely the Native American genetic component, depicted in green (Fig. 3B). Regarding maternal genetic ancestry, we identified an A2+(64) mitochondrial haplogroup, which is commonly found in Native American individuals. In addition, we estimated that the 87Sr/86Sr ratio measured on the individual COYC5 bone (0.704866) is similar to the average 87Sr/86Sr ratio (0.70465–0.70487) found in soils and rocks from the Mexico Valley (Table S9) (Pacheco-Forés et al., 2020). This indicated that the individual had spent some time in the Valley of Mexico before her death.

**Figure 3.**
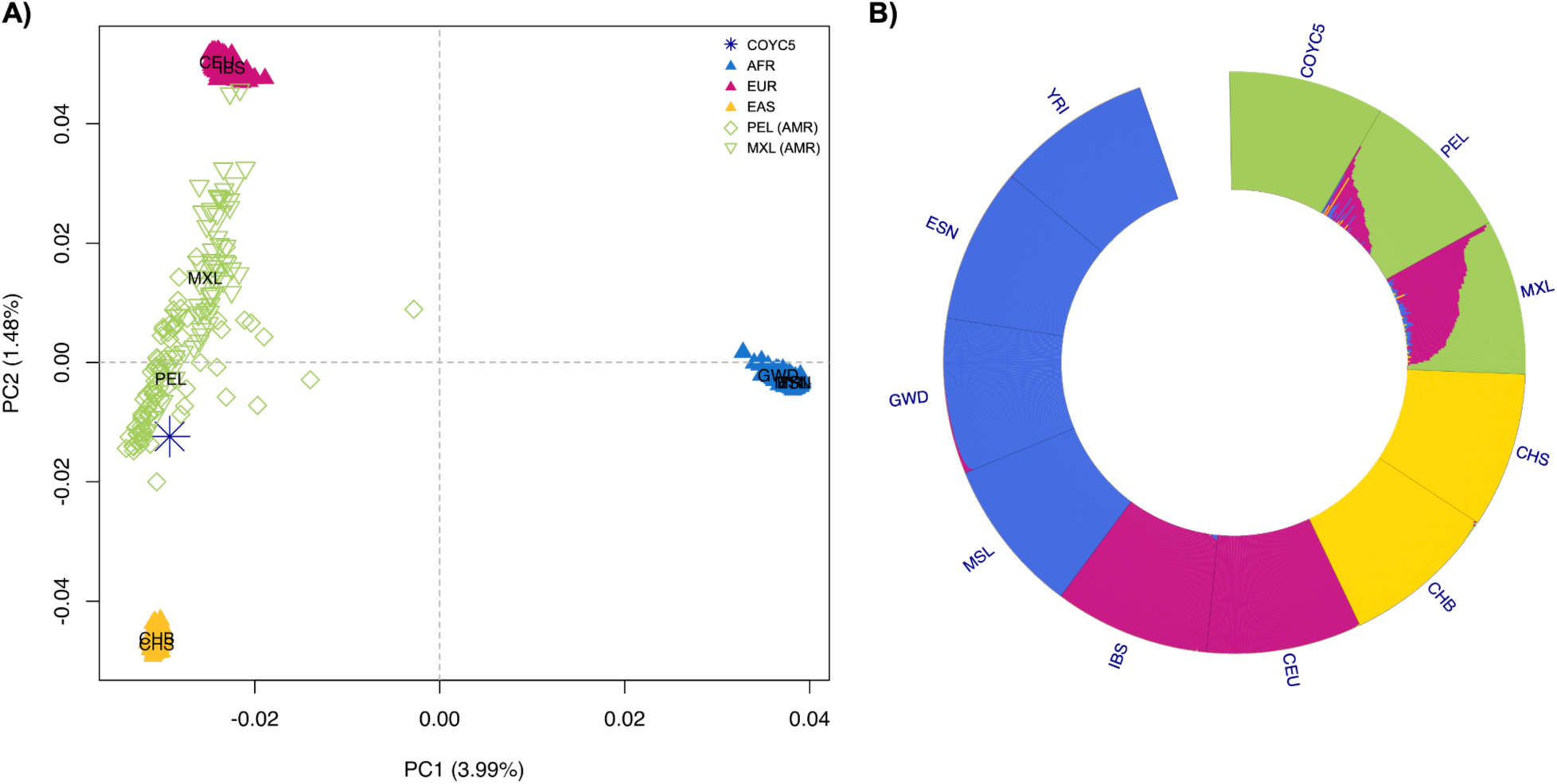
The genetic ancestry of the individual COYC5. (A) The Principal Component Analysis (PCA) of COYC5 compared to the 1000 Genomes Project reference panel. COYC5 is shown with a blue asterisk. Four super populations are represented by different colors and are labeled using a three-letter code from the 1000 Genomes Project: EUR for Europeans (IBS, CEU), EAS for East Asians (CHB), AMR for admixed populations from the Americas (MXL, PEL), SAS for South Asians (CHS), and AFR for Africans (YRI, MSL). (B) ADMIXTURE analysis of COYC5 intersected with the 1000 Genomes ancestry-informative sites (see methods). The analysis was run with k= 4 for 100 replicates, the plot shows the run with the best likelihood. Each color in the graph represents a different genetic component.

### 2.6. Presence of SPI-7 and active shufflon mechanism in COYC5

*S. enterica* pathogenicity islands (SPIs) are clusters of genes encoding virulence factors essential for host colonization and invasion (Hensel, 2004; Sia et al., 2025). We evaluated the presence and absence of 12 SPIs and the pSPCV plasmid, with a focus on the COYC5 genome (Figure 3, Table S10). To facilitate comparison across individuals, we calculated the depth of coverage for each SPI using min–max normalization (see Methods; Figures S10–S14).

Consistent with previous studies (Key et al., 2020; Wu et al., 2021; Zhou et al., 2018), SPI-6 and SPI-7 exhibited differences between ancient and modern *S. enterica* Paratyphi C genomes. The detection of the pSPCV plasmid in COYC5 confirms the genetic potential for the systemic virulence characteristic of serovar Paratyphi C, consistent with that observed in modern strains (Figure 4, Figure S14) (Liu et al., 2009).

**Figure 4.**
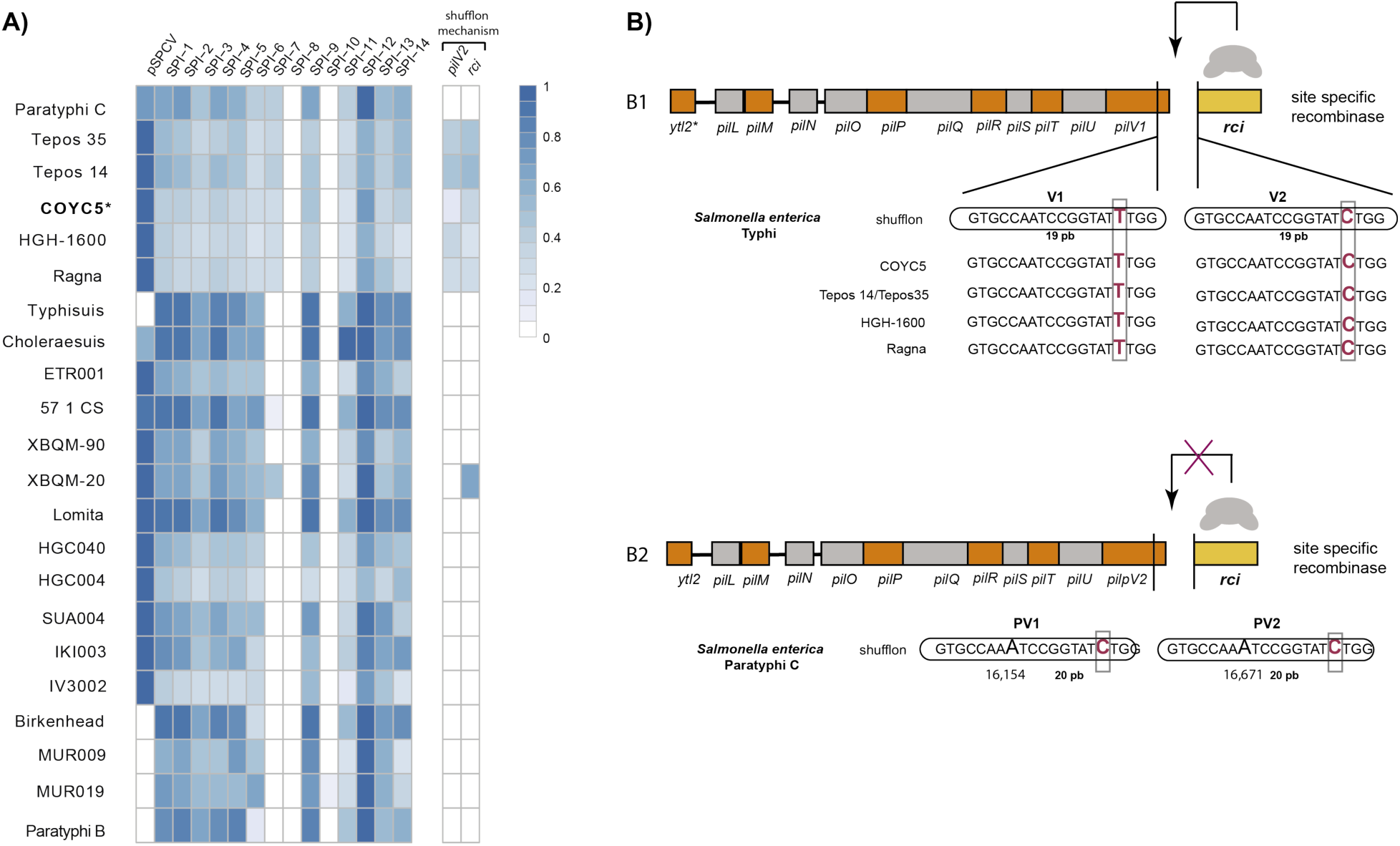
Heatmap showing the presence and absence of *Salmonella* pathogenicity islands (SPI-1 to SPI-14) and virulence factors, including the pSPCV plasmid, *pilV2*, and *rci.* The normalized depths of the SPIs and virulence factors for all analyzed individuals are shown in Figures S12 and S13. B) Comparison of the pil operons and their flanking regions (including the upstream *ytl2* gene and the downstream *rci* gene) between serovar Typhi and serovar Paratyphi C strain SGSC 2712, modified from Tam et al. (2004). (B1) In serovar Typhi, two 19-bp inverted repeats (V1 and V2), differing by a single base (underlined), serve as recognition sites for Rci, which mediates DNA inversion between them. The V1 and V2 sequences are shown for COYC5, Tepos, HGH-1600, and Ragna. (B2) In serovar Paratyphi C, each repeat contains an additional base pair (shown in larger typeface), and the shufflon appears fixed in the PV2 orientation. The nucleotide positions of the serovar Paratyphi C are depicted according to the reference sequence GenBank AY249242, where bp 1 is positioned upstream of the topoisomerase B gene (*topB*).

The COYC5 genome retained SPI-7, a major pathogenicity island carrying *pilV2* and *rci*, key genes that mediate pili diversity and virulence. Alongside Tepos 14 and Tepos 35 and the European genomes Ragna and HGH-1600, COYC5 preserves both genes (Figure 4). The *pilV2* gene encodes a pilus adhesin and the virulence-associated capsular polysaccharide (Vi antigen), which mediates phagocyte-mediated death and systemic dissemination through the IVB operon (Worley, 2023). The *rci* gene encodes a tyrosine site-specific recombinase that drives the shufflon mechanism, a DNA recombination system generating structural diversity in type IVB pili. The shufflon rearranges invertible DNA segments flanked by 19-bp inverted repeats, producing multiple pilV variants with different C-terminal regions. This enables a single genome to produce pili with variable surface properties, which affect adhesion, host interaction, auto-aggregation, and immune evasion, thereby contributing to phenotypic plasticity and adaptation (Tam et al., 2004).

The conserved presence of SPI-7 and an active shufflon in COYC5, along with the Southern Mexico (Tepos 14 and 35) and European genomes (HGH-1600 and Ragna), indicates that these ancient *S. enterica* Paratyphi C strains likely shared similar mechanisms for host infection and adaptation. This reflects a conserved virulence strategy that may have contributed to their successful systemic dissemination across different regions and time periods in Mexico and Europe.

## 3. Discussion

This study provides genomic evidence of an ancient *Salmonella enterica* Paratyphi C circulating in Mexico City between 1800 and 1899, thereby extending the known geographic and temporal range of the pathogen in Mexico. During the 19th century, typhoid fever became a recurring disease among the Mexican population, prompting physicians to focus on the disease and propose a specific diagnosis (Jiménez, 1846; Jiménez, 1982). Historical accounts describe dense settlement and poor sanitation during 1800-1850 CE (Márquez Morfín, 1991)—conditions that would have facilitated the occurrence of epidemics, such as smallpox, measles, and typhus, among others. Burial practices at “La Conchita”, including the application of lime to some graves, further suggest concern over the contagion of infectious diseases (Salas Contreras, 2007). Other pathogens have been identified at the same site through paleogenomics, such as *T. forsythia* (Bravo-Lopez et al., 2020) and Human Parvovirus B19 (Guzmán-Solís et al., 2021), underscoring that multiple infections were part of the disease environment during the 19th century (1800-1899 CE) in Mexico.

The *S. enterica* COYC5 genome from “La Conchita” groups closely with Teposcolula genomes from southern Mexico (Tepos), and the medieval European strains fall basal to this cluster in the phylogeny. This placement supports a European introduction of *S. enterica* Paratyphi C during colonization, followed by local diversification within Mexico.

The divergence estimation times obtained indicate that the *S. enterica* Paratyphi C lineage introduced into colonial Mexico originated from a deeply divergent European ancestor ∼1,400 yBP. Following its arrival, the lineage underwent local diversification, with the split between the COYC5 and southern Mexico genomes (∼516 yBP) coinciding with the early colonial period. Consistent with this pattern, the split between the F1691-1810 individual and the Teposcolula genomes, estimated at 755–541 yBP (de-Dios et al., 2021), suggests that the Paratyphi C strains present in Mexico descended from a European ancestor that had already diversified centuries before European colonization (∼early 1500s). This reinforces the hypothesis that the pathogen’s introduction into the Americas involved a lineage that was already established and genetically diversified in Europe well before it was introduced to the Americas.

Across these genomes, the presence of SPI-7 and shufflon-related genes (*pilV2* and *rci)* highlights a striking conservation of virulence mechanisms across time and geography. The *rci* recombinase, which mediates pilus variation and bacterial self-association (Tam et al., 2004), may have contributed to the pathogen’s ability to colonize human hosts effectively. This conservation suggests that *S. enterica* Paratyphi C retained a robust pathogenic potential in Mexico centuries after its introduction to the Americas.

In parallel, genetic variation in human populations has also shaped susceptibility and resistance to *S. enterica.* In 14th-century Germany, individuals infected with *S. enterica* Paratyphi C exhibited elevated frequencies of the HLA-DRB1*03:01 allele, a variant associated with an increased risk of enteric fever, compared to both contemporaneous and modern populations (Haller, Bonczarowska, et al., 2021). By contrast, HLA-DR4 alleles—linked to greater resistance against *S. enterica*—have been found enriched in both present-day and ancient Maya populations (Barquera et al., 2024). These immunogenetic patterns suggest that host–pathogen interactions with *S. enterica* Paratyphi C could have differentially impacted European and Mexican populations. Ongoing work aims to recover and characterize HLA alleles from the COYC5 individual, which may provide direct insights into host susceptibility and immune responses in 19th-century Mexico.

At the same time, ecological and social conditions must also be considered: While the Teposcolula genomes from the 16th century were linked to a major epidemic context, the “La Conchita” genome from the 19th century derives from an urban burial not associated with a known outbreak. This temporal and contextual gap raises the possibility that *S. enterica* Paratyphi C, initially an epidemic pathogen in early colonial Mexico, may have later persisted as an endemic infection under different demographic and environmental conditions. Alternatively, this bacterium may not have been the causative agent of the *Cocoliztli* epidemic, but rather found in association with victims who died from other yet-undetected pathogens, including RNA hemorrhagic viruses, which have a better match to the described symptoms, as proposed by others (Acuña-Soto et al., 2002). Exploring additional epidemic and non-epidemic contexts, as well as characterization of the host genetics, are essential for testing different hypotheses.

Lastly, this study underscores the methodological advancements in capturing and enriching ancient pathogen genomes. Capture-enrichment has become a cornerstone of ancient DNA research, whether using RNA probes (Arbor myBaits), DNA baits (Twist Biosciences), or marker-based parallel capture (Kocher et al., 2025). Here, we demonstrate that an in-house whole-genome RNA bait protocol (Carpenter et al., 2013), complemented by deep sequencing, achieved ∼11-fold coverage of an ancient *S. enterica* Paratyphi C genome, highlighting a cost-effective alternative to commercial kits and demonstrating the utility of tailored enrichment approaches for exploring local pathogen diversity.

Looking ahead, expanding the paleogenomic record of *S. enterica* Paratyphi C will be key to reconstructing its evolutionary history in the Americas and beyond. Higher-coverage sequencing of genomes, such as F1691-1810, will help resolve current phylogenetic uncertainties. Meanwhile, broader sampling from both epidemic and non-epidemic contexts will clarify how the pathogen adapts to different ecological settings. Such efforts will move us closer to understanding how *S. enterica* Paratyphi C persisted in the past and how its interactions with human populations shaped its long-term trajectory.

## 4. Conclusion

The presence of *S. enterica* Paratyphi C at “La Conchita” in an individual from 1800-1899 provides direct evidence that this pathogen affected individuals beyond the Teposcolula-Yucundaa site in southern Mexico during the 1545–1548 *Cocoliztli* epidemic, revealing a broader geographic and temporal footprint of this historically infectious disease. The ecological and social context of “La Conchita” likely allowed *S. enterica* to diversify and establish itself as a persistent pathogen, as evidenced by its presence in the COYC5 individual, who also presented osteological indicators of physical stress, reflecting the health vulnerabilities of individuals exposed to this pathogen.

Our analyses indicate that *S. enterica* Paratyphi C, introduced from Europe, diversified locally within Mexico, reflecting microbial exchanges shaped by colonial contact. This is observed in the phylogenetic placement of COYC5 alongside Teposcolula genomes, nested within a clade derived from European strains, which supports a model of transatlantic introduction, as proposed by de-Dios et al. (2021). The conserved presence of key virulence loci, including SPI-7 and the shufflon system, demonstrates the persistent pathogenic potential of *S. enterica* Paratyphi C for over two hundred years. These results raise the question of whether host genetic diversity primarily influences the severity and spread of paratyphoid fever across different temporal and environmental contexts in Mexico. Expanding genomic datasets of both ancient and modern *S. enterica* from Mexico will be essential for refining divergence estimates, resolving phylogenetic uncertainties, and understanding the true pathogenic potential, as well as how *S. enterica* Paratyphi C adapted to diverse hosts and environments over time.

## 5. Materials and Methods

### 5.1. Samples and DNA extraction

Tooth samples were obtained from the Temple of the Immaculate Conception, “La Conchita”, Mexico City, Mexico (n = 7 teeth) (Table S1). These individuals were originally part of two independent studies: Bravo-Lopez et al. (2020) and Guzmán-Solís et al. (2021). Permits to carry out DNA analyses with these samples were granted by the Archaeology Council of the Instituto de Antropología e Historia (INAH) of Mexico with permit number 401.1 S.3-2020/1310.

The only skeletal element recovered from *S. enterica*-positive COYC5 was a skull that showed no osteological evidence of a specific pathological condition. The skull belonged to a female of approximately 14 years of age (Figure S1). The remains correspond to a secondary deposition within a simple burial pit (Figure S2). Evidence of physiological stress was minimal, limited to slight enamel hypoplasia, mild alveolar resorption, and periodontal inflammation. Additionally, subtle lipping of the occipital condyles was observed, likely reflecting axial loading stress associated with physically demanding activity.

The tooth sample from COYC5 was processed in the aDNA facilities of the International Laboratory for Human Genome Research at UNAM. The dental piece was cut at the enamel-dentin junction, and approximately 200 mg of powder drilled from the tooth roots was used to perform aDNA extraction according to Dabney et al. (2013).

### 5.2. Sequencing library preparation and sequence data quality control

The resulting aDNA extracts were transformed into double-stranded Illumina libraries and shotgun sequenced on an Illumina NextSeq550 (2 × 75 cycles) at LANGEBIO’s genomics core facility (National Laboratory of Genomics for Biodiversity, Irapuato, Guanajuato). Additionally, three negative controls, carried along with DNA extraction, library preparation, and PCR indexing, were sequenced. Processing of fastq files (clipping adapters and merging sequence pairs) was performed using AdapterRemoval v2 (Schubert et al., 2016). Reads were collapsed with a minimum overlap of 11 bp, and only merged sequences ≥30 bp in length and with a base quality of ≥33 were retained for downstream analyses. We mapped filtered reads against the human reference genome (version GRCh38), using the ‘*BWA aln’* option with (flags −l500 −t 16 −q 25). We selected the unmapped reads and performed metagenomic screening of the seven individuals for human pathogens through *Kraken2* (Wood et al., 2019) using the database PlusPFP (version 10/9/2023) composed of bacterial, archaeal, viral, plasmid, human, UniVec_Core, protozoa, and fungi genomes (available at https://benlangmead.github.io/aws-indexes/k2) (Table S2). The visualization of the relative abundance of taxonomic groups was performed with the *phyloseq* library (McMurdie & Holmes, 2013).

### 5.3. Identification and reconstruction of the ancient *S. enterica* genome

#### 5.3.1. *S. enterica* authentication

The authentication of *Salmonella enterica* reads was performed through competitive mapping using a multifasta reference genome, which contained the reference genomes listed in Table S6. We used *samtools* to exclude reads flagged as repetitive (XT:A:R) or with alternative mapping positions (XA:Z), and retained only reads with a unique best alignment (X1:i:0). The resulting BAM files contained uniquely mapping non-duplicated reads. The *S. enterica*-positive sample was mapped to the *S. enterica* Paratyphi C str. RKS4594 reference (NC_012125.1) using the *BWA aln* option (with the flags −l 16 −n 0.01, −q 37). Clonal duplicates were removed using *samtools rmdup* (Li & Durbin, 2009). Average depth of coverage per reference (chromosome and plasmid) was calculated using *samtools depth -a -r <REFERENCE>* to account for collapsed reads and uneven coverage. Damage and fragmentation patterns were assessed using *mapDamage2* (Ginolhac et al., 2011) (Figure S3).

#### 5.3.2. In-house capture-enrichment

We performed the WISC capture strategy as described by Carpenter et al. (2013). We constructed biotinylated RNA probes transcribed from the *Salmonella enterica* serovar Typhimurium strain SO2, isolated from a Mexican individual and previously sequenced by Silva et al. (2016). To evaluate the efficiency of the capture library, we first performed a low-depth sequencing run. After confirming successful enrichment, we proceeded with high-depth sequencing of both the pre-capture and capture libraries on an Illumina NextSeq 550 (2 × 75 cycles). The evaluation of capture-enrichment efficiency is presented in Table S5. On-target and off-target reads were determined for each library by comparing the number of uniquely mapped reads to the *S. enterica* Paratyphi C reference genome against the total number of sequenced reads. The statistical significance of enrichment after capture was assessed using a chi-squared test for independence. All calculations and statistical analyses were performed in RStudio (R version 2024.04.2+764), using the *chisq.test()* function on a 2×2 contingency table of on- and off-target reads before and after capture.

### 5.4. *S. enterica* phylogenetic analysis

For the phylogenetic placement of the ancient genomes, we included 221 modern *S. enterica* genome assemblies from EnteroBase (Zhou et al., 2019) and 14 previously reported ancient *S. enterica* genomes (Table S6 and S7).

Each modern genome was split into k-mers of 100 bases with a step size of 1 base and mapped against the *S. enterica* Paratyphi C RKS4594 reference following the same stringent criteria used for the ancient genomes. In addition, we included two previously published 16th-century Mexican genomes (Tepos), seven from Eurasia, and one medieval Norwegian (Ragna) genome using the publicly available raw reads (Table S7) (Key et al., 2020; Vågene et al., 2018; Zhou et al., 2018). An alignment of all variable sites was constructed for all modern and ancient genomes using the tool *Multivcfanalyzer* version 0.87-alpha (Bos et al., 2014). We excluded repetitive and highly conserved regions of the *S. enterica* Paratyphi C RKS4594 reference from SNP calling to avoid spurious read mapping (Vågene et al., 2018). Genotyping was carried out using the *UnifiedGenotyper* from GATK v3.5 (McKenna et al., 2010), with the *EMIT_ALL_SITES* option enabled to report all positions. To avoid spurious SNP calls, every site reported per genome had to have at least 5-fold coverage and a genotype support of at least 90%. We define the core genome of all alignments by using only sites shared across all genomes (complete deletion), which required removing ancient genomes with a genome-wide coverage below three-fold to avoid excessive loss of positions, resulting in an alignment with an overall 93,590 SNPs. A maximum-likelihood tree was then constructed based on this alignment using RAxML with the GTRCAT model and 1000 bootstraps (Figure 3).

### 5.5. Bayesian phylogenetic reconstruction

To investigate whether the SNP alignment contained a temporal signal, we used *TempEst* v1.5.3 with the maximum-likelihood tree generated by RAxML and the associated sample dates (Figure S7). *TempEst* indicated sufficient temporal structure to support molecular clock analyses. To further assess the strength of this signal, we performed a linear regression of root-to-tip genetic distance against sampling date using the *lm()* function in R, based on the distance and date values output by *TempEst*.

Consequently, a Bayesian time-scaled tree was created with BEAST v2.6.3 following previous *Salmonella* phylogenetic reconstructions (Figure S8) (Key et al., 2020; Zhou et al., 2018), BEAST analysis was performed employing the GTR+G substitution model for nucleotides with six rate categories, applying the Bayesian skyline coalescent model, and using a strict molecular clock, given that *TempEst* analysis above demonstrated that the data has a linear trend with small residuals values, -0.02 to 0.05 (Rambaut et al., 2016). Samples’ dates in Before Present format were incorporated into the analysis. To integrate samples beyond 1950, the date of the most recent sample (*S. enterica* Paratyphi B, a modern sample from 2015) was used as year 0, rather than 1950. Then, each sample date was adjusted to years before 2015 to run BEAST. Once the Bayesian tree was obtained, the age and time range estimates for every node were re-adjusted to years Before Present, applying an offset of *-65* years (the difference between 2015 and 1950). The BEAST run was performed for one million MCMC generations, sampling every 1,000 generations. Subsequently, a Maximum Clade Credibility (MCC) tree was generated with *TreeAnnotator*, another BEAST utility, discarding the first 10% of trees as burn-in. The output log file was analyzed using *Tracer* v1.7.2, which confirmed effective sample sizes (ESS) greater than 200 for all likelihood estimates.

### 5.6. Phylogenetic placement

We performed phylogenetic placement to incorporate four low-coverage samples into the maximum-likelihood phylogenetic tree (Figure S9). Phylogenetic placement requires variant calls for the samples to be placed. To obtain these variants, a joint variant call was performed using GATK v4.3 on all BAM files of the samples within the tree, with the reference genome used as the reference for variant calling and a ploidy of one assumed. Variants were then used with *pathPhynder* v10.1093/molbev/msac017 (Martiniano et al., 2022) to perform phylogenetic placement using default parameters. The numeric labels appended to the names of phylogenetically placed samples represent the placement scores generated by *pathPhynder*. Each label consists of four values separated by hyphens, corresponding to the number of supporting markers above the branch, conflicting markers above the branch, supporting markers on the branch, and conflicting markers on the branch. A placement is considered reliable when the values in the first and third positions are higher than those in the second and fourth. Conversely, if the second and fourth values exceed the first and third, the placement is not reliable. The phylogenetic tree, including the placed samples, was visualized using *FigTree* (http://tree.bio.ed.ac.uk/software/figtree). The four low-coverage genomes—F1691, OBP001, XBQM-7, and XBQM-64—were successfully incorporated into the known tree topology, and all showed sufficient support scores for reliable placement (Supplementary Figure 7).

### 5.7. Pairwise genetic distances

We calculated pairwise maximum likelihood genetic distances between the COYC5 and the samples that form the C lineage using RAxML v8.2 (Stamatakis, 2014) with the following parameters -model GTR+G --seed 2 --threads 9 --bs-trees autoMRE. The resulting distances were visualized as a symmetric matrix and plotted as a heatmap (Figure S6) using *Python* v3.8.

### 5.8. Absence and presence of genes

We constructed a multifasta genome that included SPI-1 to SPI-14, encompassing the virulence genes within these regions, based on *Salmonella enterica* Typhi CT18 (NC_003198). The SPI’s sequences were obtained from the PAIDB database (Yoon et al., 2015). We aligned against this reference the unmapped fastq files to the human reference genome using *BWA aln* with the flags −l 1024, −n 0.03, and−q 0. The rationale is that when BWA identifies a read that maps to more than one location, it randomly selects one and assigns it a 0-mapping quality, so when filtering by quality one, it would disregard these reads, which in turn could increase the chances of a gene with paralogous sequences in the genome being considered as absent. Coverage at the gene level was calculated using *BEDTools* (Quinlan & Hall, 2010). We used a min–max normalization to scale the values to the range of 0 to 1, making the data comparable across individuals (Figure S10-S14).

The following formula was used, where *min* and *max* are the minimum and maximum values in *X* given its range. So, *Xi* converts to *Yi*.

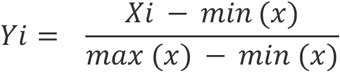

We considered genes to be absent only if they had a normalized depth value of 0. These criteria are equivalent to defining a gene as absent if its depth is zero; however, the normalization step allows us to contrast samples with very different average depth ranges across genes. The normalized depth of each gene in the ancient and modern genomes in our dataset was plotted using the *pheatmap* library in R.

### 5.9. Principal Component Analysis

Human-mapped reads from the four sequencing runs of COYC5 individual (Table S8) were used to infer the genetic ancestry of the host via principal component analysis (PCA). Genomic alignments to GRCh38 for COYC5 were intersected with genotype data from 1000 Genomes Project (1KGP) (Consortium & The 1000 Genomes Project Consortium, 2015) as a reference panel (1,560,715 SNVs). We selected 10 populations as representatives of the continental superpopulations, with African (YRI, ESN, GWD, MSL), European (IBS, CEU), East Asian (CHB, CHS), and Native American (MXL, PEL) ancestry.

From the 1KGP dataset, we extracted SNVs present on the Multi-Ethnic Genotyping Array (MEGA) to retain ancestry-informative markers (Bien et al., 2016), and applied a genotype missingness filter (--geno 0.1) using *PLINK* (Purcell et al., 2007), resulting in 1,560,715 SNVs in the final reference panel. Genotypes for the ancient individual COYC5 were called at the same sites, applying a base quality filter of >30. The merged dataset included 356,311 overlapping SNVs.

To account for low coverage, pseudo-haploid genotypes were generated by randomly sampling one allele per site in both the reference panel and COYC5. Further filtering was applied using *PLINK* with a minor allele frequency threshold of 0.05 (--maf 0.05). PCA was conducted using *smartpca* from the *EIGENSOFT* package (Patterson et al., 2006; Price et al., 2006), and genetic ancestry was also assessed with *ADMIXTURE* (Alexander et al., 2009).

### 5.10. Genetic ancestry of individual COYC5

A total of 356,311 SNPs overlapping between the COYC5 ancient genome and the 1000 Genomes Project reference panel were used for ancestry analysis (see previous section for details). *ADMIXTURE* v1.3 (Alexander and Lange, 2011) was used to estimate ancestry proportions, with K values ranging from 2 to 5. Each k was replicated 100 times with different random seeds, with k4 yielding the lowest CV error. The k4 run yielding the highest log-likelihood was selected for visualization. Ancestry proportions were plotted using *AncestryPainte*r (Feng et al., 2018).

### 5.11. Mitochondrial haplogroup and sex determination

Sequencing reads were aligned to the revised Cambridge Reference Sequence (rCRS) of the human mitochondrial genome using *BWA aln* with default parameters. The resulting alignment was processed with *Schmutzi* (Renaud et al., 2015) to reconstruct a consensus mitochondrial sequence. Haplogroup assignment was performed using *Haplogrep 2* (Kloss-Brandstätter et al., 2011; Weissensteiner et al., 2016) based on this consensus sequence.

Biological sex was inferred by calculating the proportion of reads mapped to the Y chromosome (Ry) relative to the total reads mapped to both sex chromosomes, following Skoglund et al. (2013). Individuals with Ry < 0.016 were classified as XX, and those with Ry > 0.075 as XY. The genetic sex determination (XY) was consistent with the morphological assessment.

### 5.12. Radiocarbon dating

Radiocarbon dating for the COYC5 individual was performed at the Physics Institute of the National Autonomous University of Mexico (UNAM). Parietal bones were first cleaned, dried, and ground into powder before undergoing sequential chemical treatments: 0.5 M HCl digestion, followed by 0.01 M NaOH and 0.2 M HCl. Collagen was subsequently filtered using a cutoff of >30 kDa, and graphitization was carried out using an AGEIII (Ion Plus) system. The resulting graphite was analyzed for ^14C, ^13C, and ^12C isotopes using a Tandetron Mass Spectrometer (High Voltage Engineering Europa B.V.) equipped with a 1 MV accelerator. Radiocarbon ages were calibrated with the IntCal13 curve (Reimer et al., 2013) and processed using OxCal v4.2.4 (Bronk Ramsey, 1995).

To add a temporal stamp to the phylogenetic analysis, including the COYC5 individual, we performed radiocarbon dating, which yielded a bimodal calibrated distribution (Figure S5). The full 2σ (95.4%) range spans AD 1680–1940; however, based on the archaeological context, the individual’s death most likely occurred between 1800 to 1899 CE (Meraz, personal communication, 2023). Therefore, we selected the interval representing the highest probability density (67.7%), AD 1800–1940, as the most accurate for downstream analyses

Supporting this, stratigraphic analysis places COYC5 (Skull 5 from Burial 86) in a deposit dated between the 18th and late 19th centuries (ca. 1700–1900), when in-church burials ceased. The burial context is secondary, suggesting the body decomposed elsewhere and was later reinterred—likely due to repeated exhumations carried out during this period to create space for continued burials (Meraz, personal communication, 2023).

### 5.13. Sr isotope analysis

Strontium (Sr) isotopic analyses were performed using a THERMO SCIENTIFIC TRITON PLUS thermal ionization mass spectrometer (TIMS) at the Laboratorio Universitario de Geoquímica Isotópica (LUGIS), Instituto de Geofísica, UNAM. The TRITON instrument is equipped with nine adjustable Faraday collectors and five ion counters. All measurements were conducted in static mode.

Strontium samples were loaded as chlorides onto double rhenium filaments and measured as metallic ions. For each analytical run, 60 isotopic ratios were recorded. The integrated software identifies outliers based on signal stability during data acquisition. Reported values (1sd = ±1σ abs) represent the analytical uncertainty for the last two digits; 1SE(M) = 1σ abs /√n. All Sr isotopic ratios were corrected for mass fractionation by normalization to ⁸⁶Sr/⁸⁸Sr = 0.1194.

LUGIS standard values for NBS 987 (Sr) during the period of analysis were ⁸⁷Sr/⁸⁶Sr = 0.710256 ± 13 (±1σ abs, n = 99). The analytical blank obtained during the measurement period was 2.88 ng Sr (chemical blank). Strontium concentrations were determined by Isotope Dilution using an ⁸⁴Sr tracer.

## Supporting information

Supplementary Material, and will be used for the link to the file on the preprint site

Supplementary Tables, and will be used for the link to the file on the preprint site

## Acknowledgments

This study was supported by the Human Frontier Science Project grant (number RGY0075/2019). MAA’s work is supported by SECIHTI’s grant CF-2023-G-957. AJV was supported by Carlsbergfondet Semper Ardens grant no. CF18-1109.

## References

1. Acuna-Soto, R., Stahle, D. W., Cleaveland, M. K., & Therrell, M. D. (2002). Megadrought and megadeath in 16th century Mexico. Emerging infectious diseases, 8(4), 360–362.

2. Alexander, D. H., Novembre, J., & Lange, K. (2009). Fast model-based estimation of ancestry in unrelated individuals. Genome research, 19(9), 1655–1664. 10.1101/gr.094052.109.

3. Alexander, D. H., & Lange, K. (2011). Enhancements to the ADMIXTURE algorithm for individual ancestry estimation. BMC Bioinformatics, 12, 246. 10.1186/1471-2105-12-246.

4. Als, D., Radhakrishnan, A., Arora, P., Gaffey, M. F., Campisi, S., Velummailum, R., Zareef, F., & Bhutta, Z. A. (2018). Global trends in typhoidal salmonellosis: A systematic review. The American Journal of Tropical Medicine and Hygiene, 99(3_Suppl), 10–19.

5. Barquera, R., Del Castillo-Chávez, O., Nägele, K., Pérez-Ramallo, P., Hernández-Zaragoza, D. I., Szolek, A., Rohrlach, A. B., Librado, P., Childebayeva, A., Bianco, R. A., Penman, B. S., Acuña-Alonzo, V., Lucas, M., Lara-Riegos, J. C., Moo-Mezeta, M. E., Torres-Romero, J. C., Roberts, P., Kohlbacher, O., Warinner, C., & Krause, J. (2024). Ancient genomes reveal insights into ritual life at Chichén Itzá. Nature, 630(8018), 912–919.

6. Bien, S. A., Wojcik, G. L., Zubair, N., Gignoux, C. R., Martin, A. R., Kocarnik, J. M., Martin, L. W., Buyske, S., Haessler, J., Walker, R. W., Cheng, I., Graff, M., Xia, L., Franceschini, N., Matise, T., James, R., Hindorff, L., Le Marchand, L., North, K. E., … PAGE Study. (2016). Strategies for enriching variant coverage in candidate disease loci on a multiethnic genotyping array. PloS One, 11(12), e0167758.

7. Bos, K. I., Harkins, K. M., Herbig, A., Coscolla, M., Weber, N., Comas, I., Forrest, S. A., Bryant, J. M., Harris, S. R., Schuenemann, V. J., Campbell, T. J., Majander, K., Wilbur, A. K., Guichon, R. A., Wolfe Steadman, D. L., Cook, D. C., Niemann, S., Behr, M. A., Zumarraga, M., … Krause, J. (2014). Pre-Columbian mycobacterial genomes reveal seals as a source of New World human tuberculosis. Nature, 514(7523), 494–497.

8. Bravo-Lopez M., Villa-Islas V., Rocha Arriaga C., Villaseñor-Altamirano AB., Guzmán-Solís ASandoval-Velasco MWesp JKAlcantara KLópez-Corral AGómez-Valdés JMejía EHerrera A., Meraz-Moreno A-. Moreno-Cabrera M de la L., Moreno-Estrada A., Nieves-Colón MA., Olvera J., Pérez-Pérez J., Iversen KH., Rasmussen S., Sandoval K., Zepeda G., Ávila-Arcos MC. (2020) Paleogenomic insights into the red complex bacteria tannerella forsythia in pre-hispanic and colonial individuals from Mexico Philosophical Transactions of the Royal Society of London. Series B, Biological Sciences 375:20190580.

9. Bronk Ramsey, C. (1995). Radiocarbon calibration and analysis of stratigraphy: The OxCal program. Radiocarbon, 37(2), 425–430.

10. Carpenter, M. L., Buenrostro, J. D., Valdiosera, C., Schroeder, H., Allentoft, M. E., Sikora, M., Rasmussen, M., Gravel, S., Guillén, S., Nekhrizov, G., Leshtakov, K., Dimitrova, D., Theodossiev, N., Pettener, D., Luiselli, D., Sandoval, K., Moreno-Estrada, A., Li, Y., Wang, J., … Bustamante, C. D. (2013). Pulling out the 1%: whole-genome capture for the targeted enrichment of ancient DNA sequencing libraries. The American Journal of Human Genetics, 93(5), 852–864.

11. Consortium, T. 1000 G. P., & The 1000 Genomes Project Consortium. (2015). A global reference for human genetic variation. In Nature (Vol. 526, Issue 7571, pp. 68–74). 10.1038/nature15393

12. Dabney, J., Knapp, M., Glocke, I., Gansauge, M.-T., Weihmann, A., Nickel, B., Valdiosera, C., Garcia, N., Paabo, S., Arsuaga, J.-L., & Meyer, M. (2013). Complete mitochondrial genome sequence of a Middle Pleistocene cave bear reconstructed from ultrashort DNA fragments. Proceedings of the National Academy of Sciences, 110(39), 15758–15763.

13. de-Dios, T., Carrión, P., Olalde, I., Llovera Nadal, L., Lizano, E., Pàmies, D., Marques-Bonet, T., Balloux, F., van Dorp, L., & Lalueza-Fox, C. (2021). Salmonella enterica from a soldier from the 1652 siege of Barcelona (Spain) supports historical transatlantic epidemic contacts. iScience, 24(9), 103021.

14. Feng, Q., Lu, D., & Xu, S. (2018). AncestryPainter: A graphic program for displaying ancestry composition of populations and individuals. Genomics, Proteomics & Bioinformatics, 16(5), 382–385. 10.1016/j.gpb.2018.05.002.

15. Gal-Mor, O., Boyle, E. C., & Grassl, G. A. (2014). Same species, different diseases: how and why typhoidal and non-typhoidal Salmonella enterica serovars differ. Frontiers in Microbiology, 5, 391.

16. Ginolhac, A., Rasmussen, M., Gilbert, M. T. P., Willerslev, E., & Orlando, L. (2011). mapDamage: testing for damage patterns in ancient DNA sequences. Bioinformatics, 27(15), 2153–2155.

17. Guzmán-Solís AA, Villa-Islas V, Bravo-López MJ, Sandoval-Velasco M, Wesp JK, Gómez-Valdés JA, Moreno-Cabrera ML, Meraz A, Solís-Pichardo G, Schaaf P, TenOever BR, Blanco-Melo D, Ávila Arcos MC. (2021). Ancient viral genomes reveal introduction of human pathogenic viruses into Mexico during the transatlantic slave trade. Elife, 10:e68612.

18. Haller, M., Bonczarowska, J. H., Rieger, D., Lenz, T. L., Nebel, A., & Krause-Kyora, B. (2021). Ancient DNA study in medieval Europeans shows an association between HLA-DRB1*03 and paratyphoid fever. Frontiers in Immunology, 12, 691475.

19. Haller, M., Callan, K., Susat, J., Flux, A. L., Immel, A., Franke, A., Herbig, A., Krause, J., Kupczok, A., Fouquet, G., Hummel, S., Rieger, D., Nebel, A., & Krause-Kyora, B. (2021). Mass Burial Genomics Reveals Outbreak of Enteric Paratyphoid Fever in the Late Medieval Trade City Lübeck. 10.2139/ssrn.3762113

20. Han, J., Aljahdali, N., Zhao, S., Tang, H., Harbottle, H., Hoffmann, M., Frye, J. G., & Foley, S. L. (2024). Infection biology of Salmonella enterica. Ecosal plus, 12(1), eesp00012023.

21. Hensel, M. (2004). Evolution of pathogenicity islands of Salmonella enterica. International Journal of Medical Microbiology: IJMM, 294(2-3), 95–102.

22. Hiyoshi, H., Tiffany, C. R., Bronner, D. N., & Bäumler, A. J. (2018). Typhoidal Salmonella serovars: ecological opportunity and the evolution of a new pathovar. FEMS Microbiology Reviews. 10.1093/femsre/fuy024

23. Jiménez, M. F. (1846). Apuntes para la historia de la fiebre petequial ó tabardillo, que se observa en Mexico: Memoir presented to the Sociedad Filoiátrica, en la sesion de 31 de octubre de 1844. México: Impr. de Cumplido.

24. Jiménez, Miguel Francisco, “El tabardillo” en Enrique Florescano y Elsa Malvido, Ensayos sobre la historia de las epidemias en México, Tomo II. México, IMSS, 1982.

25. Key, F. M., Posth, C., Esquivel-Gomez, L. R., Hübler, R., Spyrou, M. A., Neumann, G. U., Furtwängler, A., Sabin, S., Burri, M., Wissgott, A., Lankapalli, A. K., Vågene, Å. J., Meyer, M., Nagel, S., Tukhbatova, R., Khokhlov, A., Chizhevsky, A., Hansen, S., Belinsky, A. B., … Krause, J. (2020). Emergence of human-adapted Salmonella enterica is linked to the Neolithization process. Nature Ecology & Evolution, 4(3), 324–333.

26. Kloss-Brandstatter, A., Pacher, D., Schönherr, S., Weissensteiner, H., Binna, R., Specht, G. and Kronenberg, F. (2011) HaploGrep: a fast and reliable algorithm for automatic classification of mitochondrial DNA haplogroups. Hum. Mutat., 32, 25–32.

27. Kocher, A., Seguin-Orlando, A., Clavel, P., Louvel, G., Jonvel, R., Tzortzis, S., Signoli, M., Costedoat, C., & Orlando, L. (2025). A minimal hybridization capture system for the parallel enrichment and cost-effective detection of ancient human pathogens. In bioRxiv. 10.1101/2025.06.02.657376

28. Kumar, G., Kumar, S., Jangid, H., Dutta, J., & Shidiki, A. (2025). The rise of non-typhoidal Salmonella: an emerging global public health concern. Frontiers in Microbiology, 16, 1524287.

29. Lamichhane, B., Mawad, A. M. M., Saleh, M., Kelley, W. G., Harrington, P. J., 2nd, Lovestad, C. W., Amezcua, J., Sarhan, M. M., El Zowalaty, M. E., Ramadan, H., Morgan, M., & Helmy, Y. A. (2024). Salmonellosis: An overview of epidemiology, pathogenesis, and innovative approaches to mitigate the antimicrobial resistant infections. Antibiotics (Basel, Switzerland), 13(1). 10.3390/antibiotics13010076

30. Li, H., & Durbin, R. (2009). Fast and accurate short read alignment with Burrows–Wheeler transform. Bioinformatics, 25(14), 1754–1760.

31. Liu, W.-Q., Feng, Y., Wang, Y., Zou, Q.-H., Chen, F., Guo, J.-T., Peng, Y.-H., Jin, Y., Li, Y.-G., Hu, S.-N., Johnston, R. N., Liu, G.-R., & Liu, S.-L. (2009). Salmonella paratyphi C: genetic divergence from Salmonella choleraesuis and pathogenic convergence with Salmonella typhi. PloS One, 4(2), e4510.

32. Márquez Morfín, L. (1991). La desigualdad ante la muerte: epidemias, población y sociedad en la Ciudad de México (C. E. Lida (Ed.)) [Doctorado en Historia]. Centro de Estudios Históricos.

33. Martiniano, R., De Sanctis, B., Hallast, P., & Durbin, R. (2022). Placing ancient DNA sequences into reference phylogenies. Molecular Biology and Evolution, *39*(2). 10.1093/molbev/msac017

34. McKenna, A., Hanna, M., Banks, E., Sivachenko, A., Cibulskis, K., Kernytsky, A., Garimella, K., Altshuler, D., Gabriel, S., Daly, M., & DePristo, M. A. (2010). The Genome Analysis Toolkit: a MapReduce framework for analyzing next-generation DNA sequencing data. Genome Research, 20(9), 1297–1303.

35. McMurdie, P. J., & Holmes, S. (2013). phyloseq: An R Package for Reproducible Interactive Analysis and Graphics of Microbiome Census Data. PloS One, 8(4), e61217.

36. Neumann, G. U., Skourtanioti, E., Burri, M., Nelson, E. A., Michel, M., Hiss, A. N., McGeorge, P. J. P., Betancourt, P. P., Spyrou, M. A., Krause, J., & Stockhammer, P. W. (2022). Ancient Yersinia pestis and Salmonella enterica genomes from Bronze Age Crete. Current Biology: CB, 32(16), 3641–3649.e8.

37. Pacheco-Forés, S. I., Gordon, G. W., & Knudson, K. J. (2020). Expanding radiogenic strontium isotope baseline data for central Mexican paleomobility studies. PloS One, 15(2), e0229687.

38. Patterson, N., Price, A. L., & Reich, D. (2006). Population structure and eigen analysis. PLOS Genetics, 2(12), e190. 10.1371/journal.pgen.0020190.

39. Price, A. L., Patterson, N. J., Plenge, R. M., Weinblatt, M. E., Shadick, N. A., & Reich, D. (2006). Principal components analysis corrects for stratification in genome-wide association studies. Nature genetics, 38(8), 904–909. 10.1038/ng1847.

40. Purcell, S., Neale, B., Todd-Brown, K., Thomas, L., Ferreira, M. A., Bender, D., Maller, J., Sklar, P., de Bakker, P. I., Daly, M. J., & Sham, P. C. (2007). PLINK: a tool set for whole-genome association and population-based linkage analyses. American journal of human genetics, 81(3), 559–575. 10.1086/519795

41. Quinlan, A. R., & Hall, I. M. (2010). BEDTools: a flexible suite of utilities for comparing genomic features. Bioinformatics (Oxford, England), 26(6), 841–842.

42. Rambaut, A., Lam, T. T., Max Carvalho, L., & Pybus, O. G. (2016). Exploring the temporal structure of heterochronous sequences using TempEst (formerly Path-O-Gen). Virus Evolution, 2(1), vew007.

43. Reimer, P. J., Bard, E., Bayliss, A., Beck, J. W., Blackwell, P. G., Ramsey, C. B., … van der Plicht, J. (2013). IntCal13 and Marine13 Radiocarbon Age Calibration Curves 0–50,000 Years cal BP. Radiocarbon, 55(4), 1869–1887. doi:10.2458/azu_js_rc.55.16947.

44. Renaud, G., Slon, V., Duggan, A. T., & Kelso, J. (2015). Schmutzi: Estimation of contamination and endogenous mitochondrial consensus calling for ancient DNA. Genome Biology, 16, 224. 10.1186/s13059-015-0776-0.

45. Sabbagh, S. C., Forest, C. G., Lepage, C., Leclerc, J.-M., & Daigle, F. (2010). So similar, yet so different: uncovering distinctive features in the genomes of Salmonella enterica serovars Typhimurium and Typhi. FEMS Microbiology Letters, 305(1), 1–13.

46. Salas Contreras Carlos, Prácticas funerarias en la ex-iglesia de la Encarnación, “Antigua Biblioteca Iberoamericana”, 2007, Arqueología mexicana, No. 36, 116–134.

47. Schubert, M., Lindgreen, S., & Orlando, L. (2016). AdapterRemoval v2: rapid adapter trimming, identification, and read merging. BMC Research Notes, 9(1), 88.

48. Sia, C. M., Pearson, J. S., Howden, B. P., Williamson, D. A., & Ingle, D. J. (2025). Salmonella pathogenicity islands in the genomic era. Trends in Microbiology. 10.1016/j.tim.2025.02.007

49. Silva, C., Calva, E., Puente, J. L., Zaidi, M. B., & Vinuesa, P. (2016). Complete Genome Sequence of Salmonella enterica Serovar Typhimurium Strain SO2 (Sequence Type 302) Isolated from an Asymptomatic Child in Mexico. Genome Announcements, 4(2). 10.1128/genomeA.00253-16

50. Skoglund, P., Storå, J., Götherström, A., & Jakobsson, M. (2013). Accurate sex identification of ancient human remains using DNA shotgun sequencing. Journal of Archaeological Science, 40(12), 4477–4482. 10.1016/j.jas.2013.07.004.

51. Stamatakis, A. (2014). RAxML version 8: a tool for phylogenetic analysis and post-analysis of large phylogenies. *Bioinformatics (Oxford*, England), 30(9), 1312–1313.

52. Tam, C. K. P., Hackett, J., & Morris, C. (2004). Salmonella enterica serovar Paratyphi C carries an inactive shufflon. Infection and Immunity, 72(1), 22–28.

53. Vågene, Å. J., Herbig, A., Campana, M. G., Robles García, N. M., Warinner, C., Sabin, S., Spyrou, M. A., Andrades Valtueña, A., Huson, D., Tuross, N., Bos, K. I., & Krause, J. (2018). Salmonella enterica genomes from victims of a major sixteenth-century epidemic in Mexico. Nature Ecology & Evolution, 2(3), 520–528.

54. Wang, B. X., Butler, D. S., Hamblin, M., & Monack, D. M. (2023). One species, different diseases: the unique molecular mechanisms that underlie the pathogenesis of typhoidal Salmonella infections. Current Opinion in Microbiology, 72(102262), 102262.

55. Weissensteiner, H., Pacher, D., Kloss-Brandstätter, A., Forer, L., Specht, G., Bandelt, H. J., Kronenberg, F., Salas, A., & Schönherr, S. (2016). HaploGrep 2: mitochondrial haplogroup classification in the era of high-throughput sequencing. Nucleic acids research, 44(W1), W58–W63. 10.1093/nar/gkw233.

56. World Health Organization. (2023). Typhoid. WHO. https://www.who.int/news-room/fact-sheets/detail/typhoid.

57. World Health Organization. (2024). WHO Bacterial Priority Pathogens List, 2024: bacterial pathogens of public health importance to guide research, development and strategies to prevent and control antimicrobial resistance.

58. Worley, M.J. (2023) Salmonella bloodstream infections. Trop. Med. Infect. Dis. 8:487

59. Wood, D. E., Lu, J., & Langmead, B. (2019). Improved metagenomic analysis with Kraken 2. Genome Biology, 20(1), 257.

60. World Health Organization. (2024). WHO Bacterial Priority Pathogens List, 2024: bacterial pathogens of public health importance to guide research, development and strategies to prevent and control antimicrobial resistance.

61. Wu, X., Ning, C., Key, F. M., Andrades Valtueña, A., Lankapalli, A. K., Gao, S., Yang, X., Zhang, F., Liu, L., Nie, Z., Ma, J., Krause, J., Herbig, A., & Cui, Y. (2021). A 3,000-year-old, basal S. enterica lineage from Bronze Age Xinjiang suggests spread along the Proto-Silk Road. PLoS Pathogens, 17(9), e1009886.

62. Yoon, S. H., Park, Y.-K., & Kim, J. F. (2015). PAIDB v2.0: exploration and analysis of pathogenicity and resistance islands. Nucleic Acids Research, 43(Database issue), D624–D630.

63. Zhou, Z., Lundstrøm, I., Tran-Dien, A., Duchêne, S., Alikhan, N.-F., Sergeant, M. J., Langridge, G., Fotakis, A. K., Nair, S., Stenøien, H. K., Hamre, S. S., Casjens, S., Christophersen, A., Quince, C., Thomson, N. R., Weill, F.-X., Ho, S. Y. W., Gilbert, M. T. P., & Achtman, M. (2018). Pan-genome Analysis of Ancient and Modern Salmonella enterica Demonstrates Genomic Stability of the Invasive Para C Lineage for Millennia. Current Biology: CB, 28(15), 2420–2428.e10.

64. Zhou, Z., Alikhan, N. F., Mohamed, K., Achtman, M., Group, A. S., & Others. (2019). The user’s guide to comparative genomics with EnteroBase. Three Case Studies: Micro-Clades within Salmonella Enterica Serovar Agama, Ancient and Modern Populations of Yersinia Pestis, and Core Genomic Diversity of All Escherichia. bioRxiv, 10(613554), 18.

