## Supplementary Material, and will be used for the link to the file on the preprint site for "Ancient genomic insights into *Salmonella enterica* Paratyphi C from Central Mexico"

1. Burial and osteological description of the COYC5 individual

Stratigraphic analysis of Burial 86, where COYC5 was found, places the deposit between the 18th and late 19th centuries—the period when in-church burials ceased. Together with radiocarbon results, this suggests the individual most likely died in the 19th century (1800–1899 CE). The burial context is secondary, indicating that decomposition occurred elsewhere (Meraz, personal communication, 2023). This reflects a practice necessitated by frequent exhumations to accommodate new interments. Morphological sex estimation of COYC5 was conducted using Morphopasse (Klales 2020), which allows the analyst to score cranial features such as the nuchal crest, mastoid process, supraorbital margin, and glabellar projection. Based on the slight degree of expression and minimal development of these traits, the results indicate a very high probability that the individual was female.

Age at death was estimated from the cranium (mandible absent) using the degree of spheno-occipital suture closure (Lottering et al. 2015) and dental eruption patterns (AlQahtani et al. 2010). The spheno-occipital synchondrosis showed an advanced state of fusion, with more than half of its length fused. The permanent dentition was fully erupted, although the third molar was still completing root apex formation. Taken together, these observations suggest that the individual was between 14 and 17 years old at the time of death, placing her within the adolescent age category (13–20 years) defined by White and Folkens (2005).

Pathological assessment followed the guidelines of Ortner (2003). Overall, only minor indicators of stress or disease were present: up to two lines of enamel hypoplasia on the canines and premolars, mild alveolar resorption, and signs of periodontal inflammation. However, these features do not indicate a specific pathological condition (Goodman and Martin 2002; Larsen 2018). In addition, slight lipping of the occipital condyles was observed, which may reflect axial load stress associated with physically demanding activity (Knüsel et al. 1997; Villotte and Knüsel 2013).

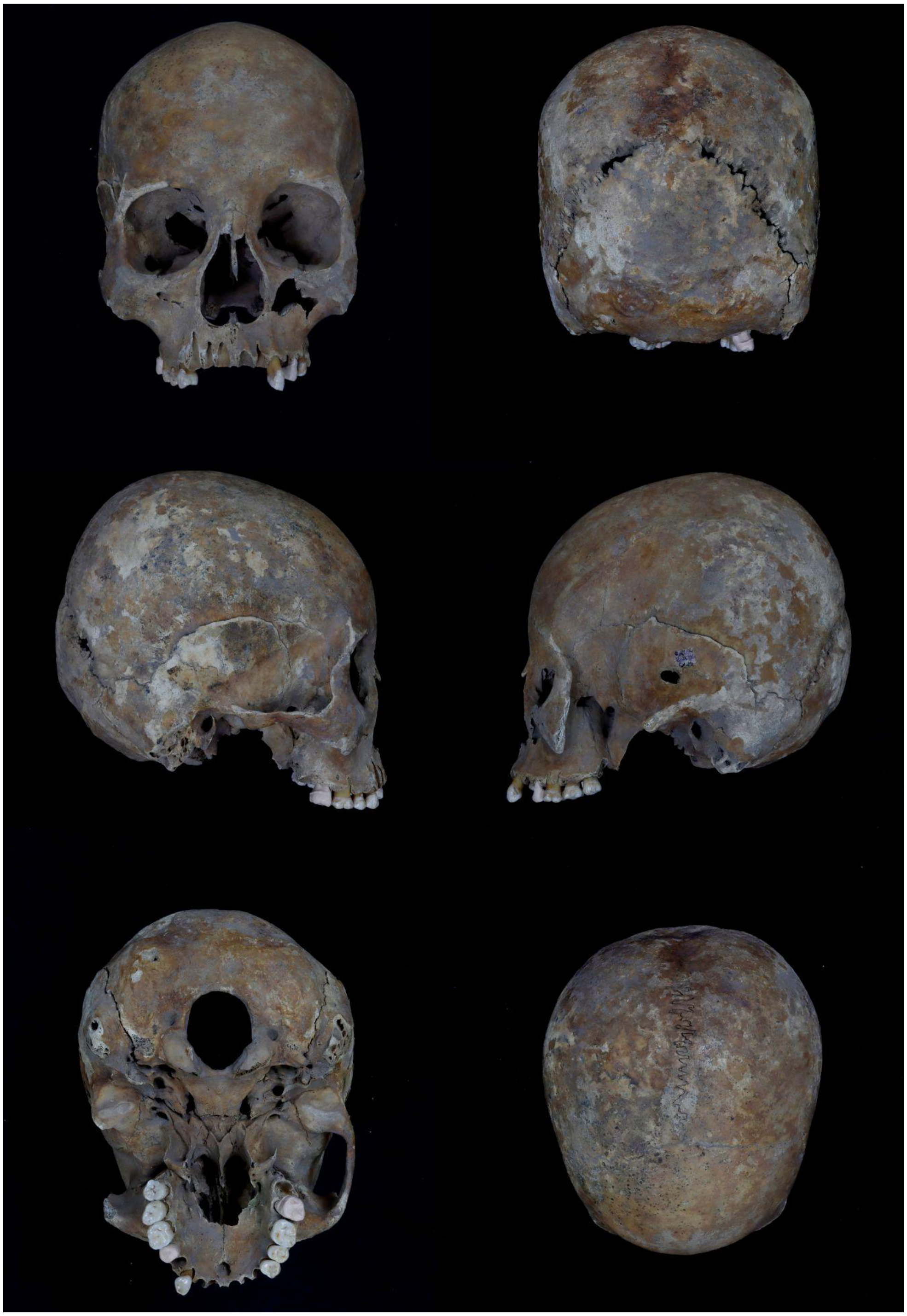

Figure S1. Lateral and superior view of the COYC5 individual skull. Credit by Jorge Gómez Valdes.

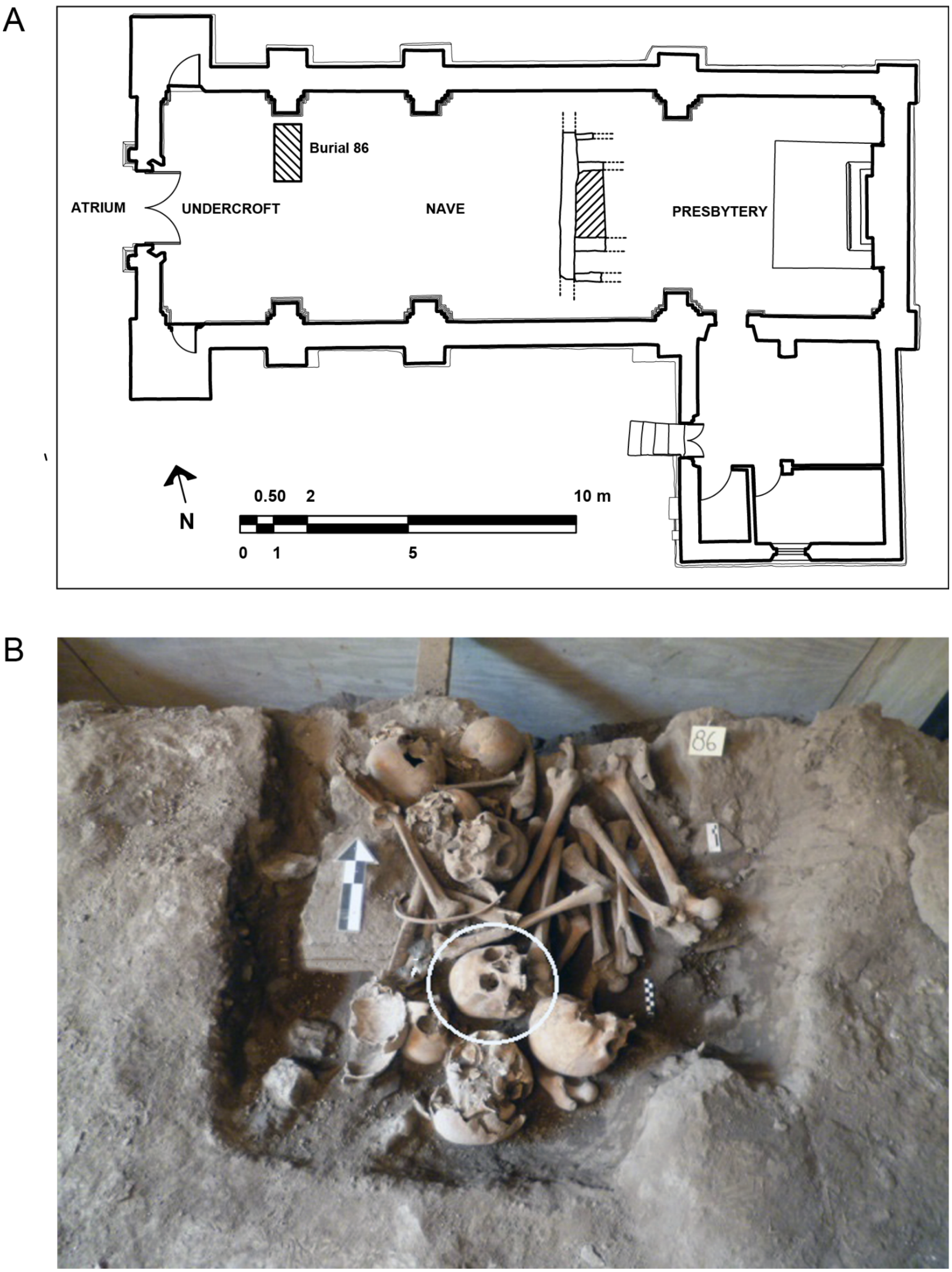


Figure S2. Location of the COYC5 human remains. A. Site plan of the Temple of the Immaculate Conception, “La Conchita”, showing the location of burial 86. Digitized by A. Meraz. B. Location of COYC5 from Burial 86, indicated by a white circle. Source: RATCC, DSA-INAH. Photo by Octavio Vargas Carranza, modified by A. Meraz.

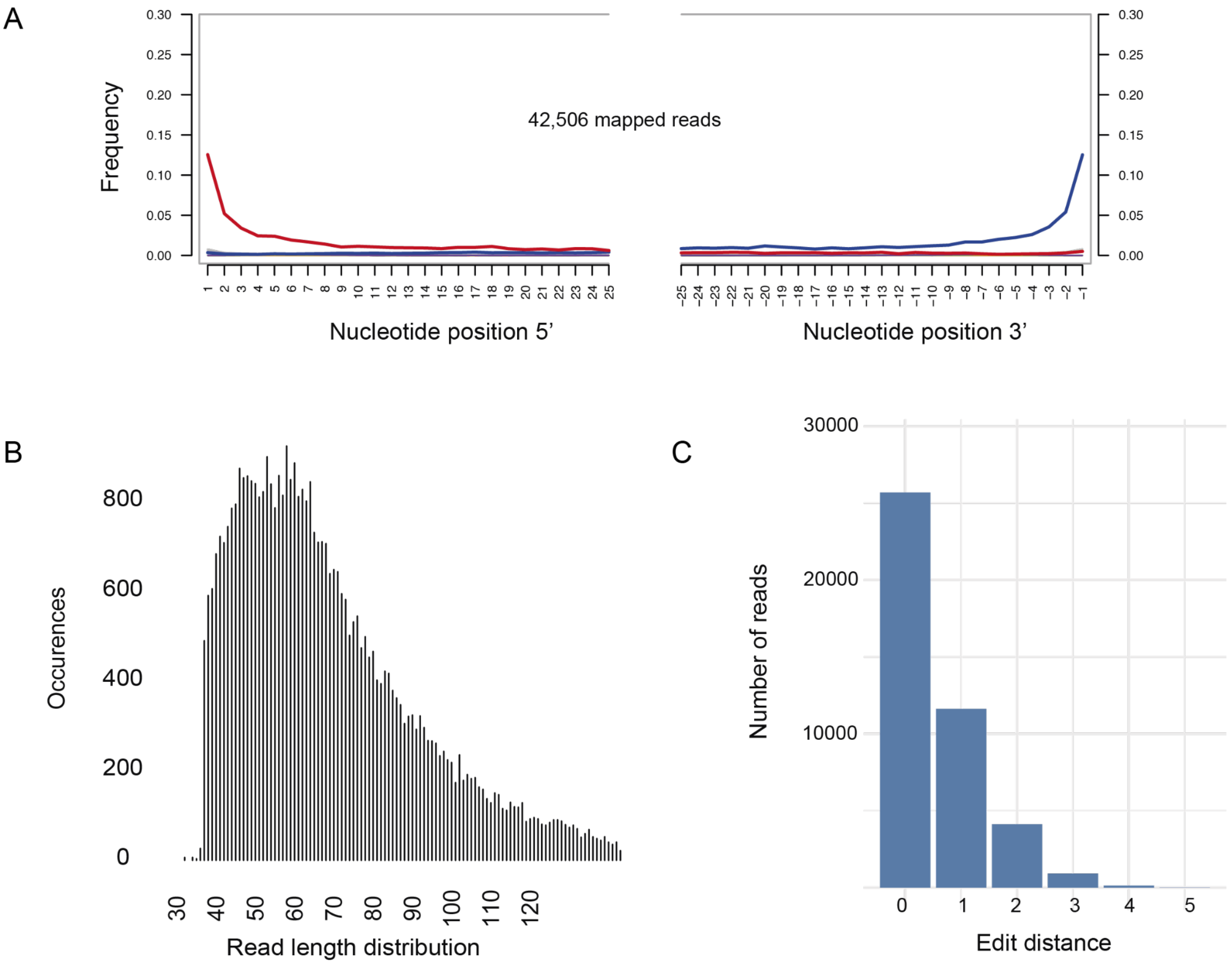


Figure S3. Authenticity of ancient DNA (aDNA) features of *Salmonella enterica* Paratyphi C in the COYC5 individual. (A) Deamination patterns showing the proportion of aDNA damage as C→T and G→A substitutions at the first position of the 5′ and 3′ ends. B) The mean read length (bp) is also indicated; the y-axis represents the number of occurrences. (C) Edit distance distribution of reads mapped to *S. enterica* Paratyphi C str. RKS4594 reference genome (NC_012125.1).

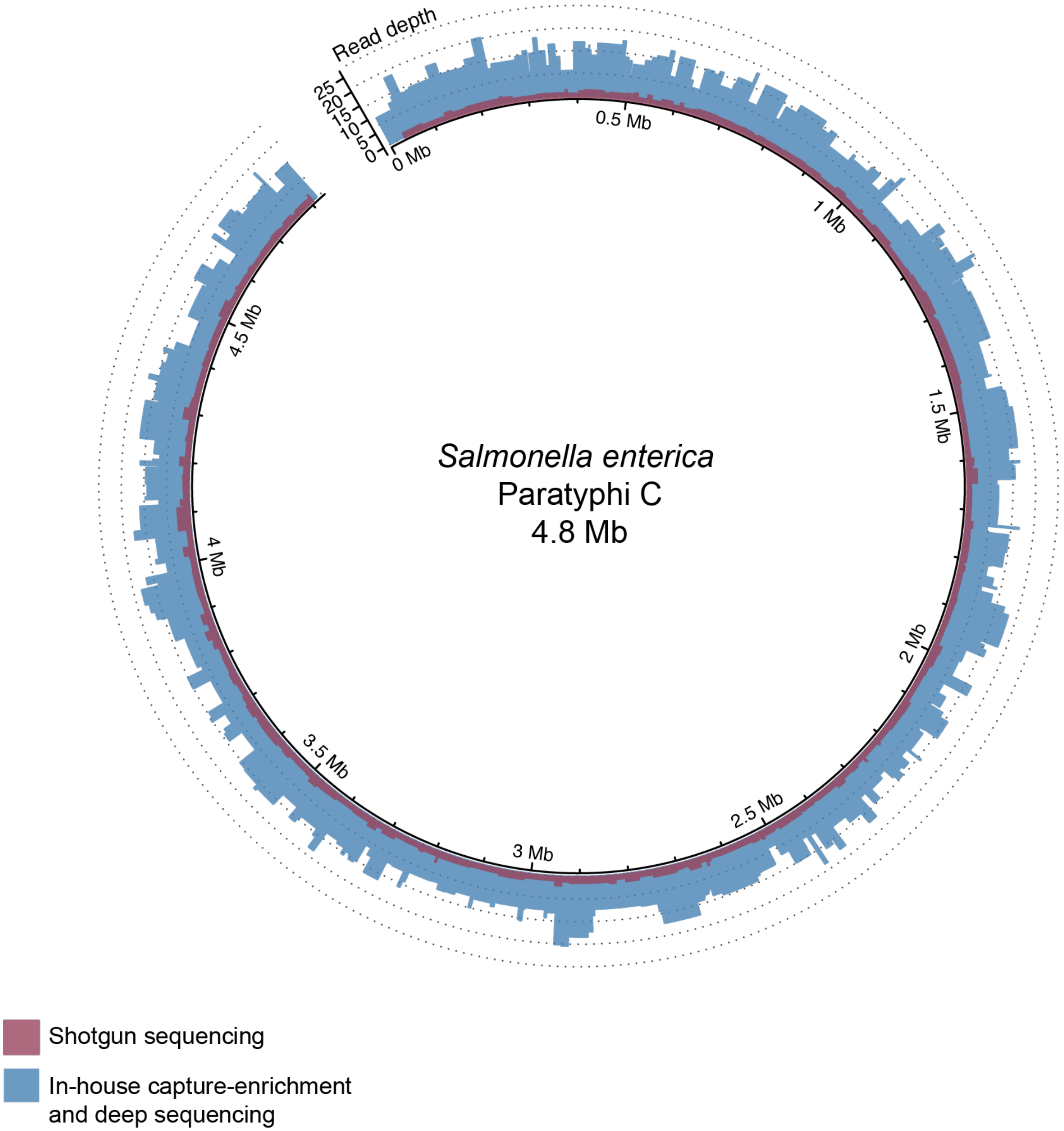


Figure S4. Circos plot showing the genome-wide distribution of reads. Shotgun sequencing data are shown in pink, while merged data from in-house capture enrichment and deep sequencing are also displayed. The plot was generated using the R package *circlize* (Gu et al. 2014).

| Name<br>Show all<br>Show structure | | Unmodelled (BC/AD) | | | | | | Select<br>All<br>Visible | Page break |
|---|---|---|---|---|---|---|---|---|---|
| | | from | to | % | from | to | % | | |
| R_Date LEMA 1671.1.1 | | 1693 | 1919 | 68.3 | 1680 | 1940 | 95.4 | ☑ 2 | ☐ |
| | | Warning! Date may extend out of range - 117+/-30BP<br>Warning! Date probably out of range - 117+/-30BP | | | | | | | |

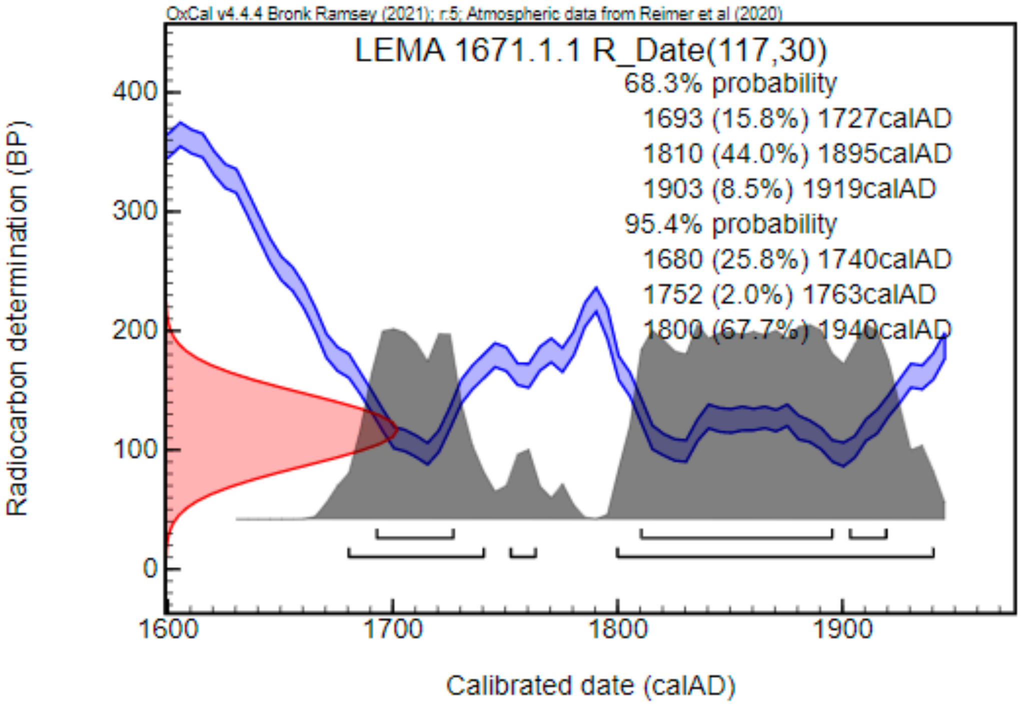


Figure S5. Radiocarbon dating of COYC5. The left axis represents radiocarbon age in years before present (BP), while the bottom axis shows calibrated calendar years derived from tree-ring data. The blue curves indicate radiocarbon measurements from the calibration dataset (±1 standard deviation), and the red curve on the left denotes the measured radiocarbon concentration of the COYC5 individual.

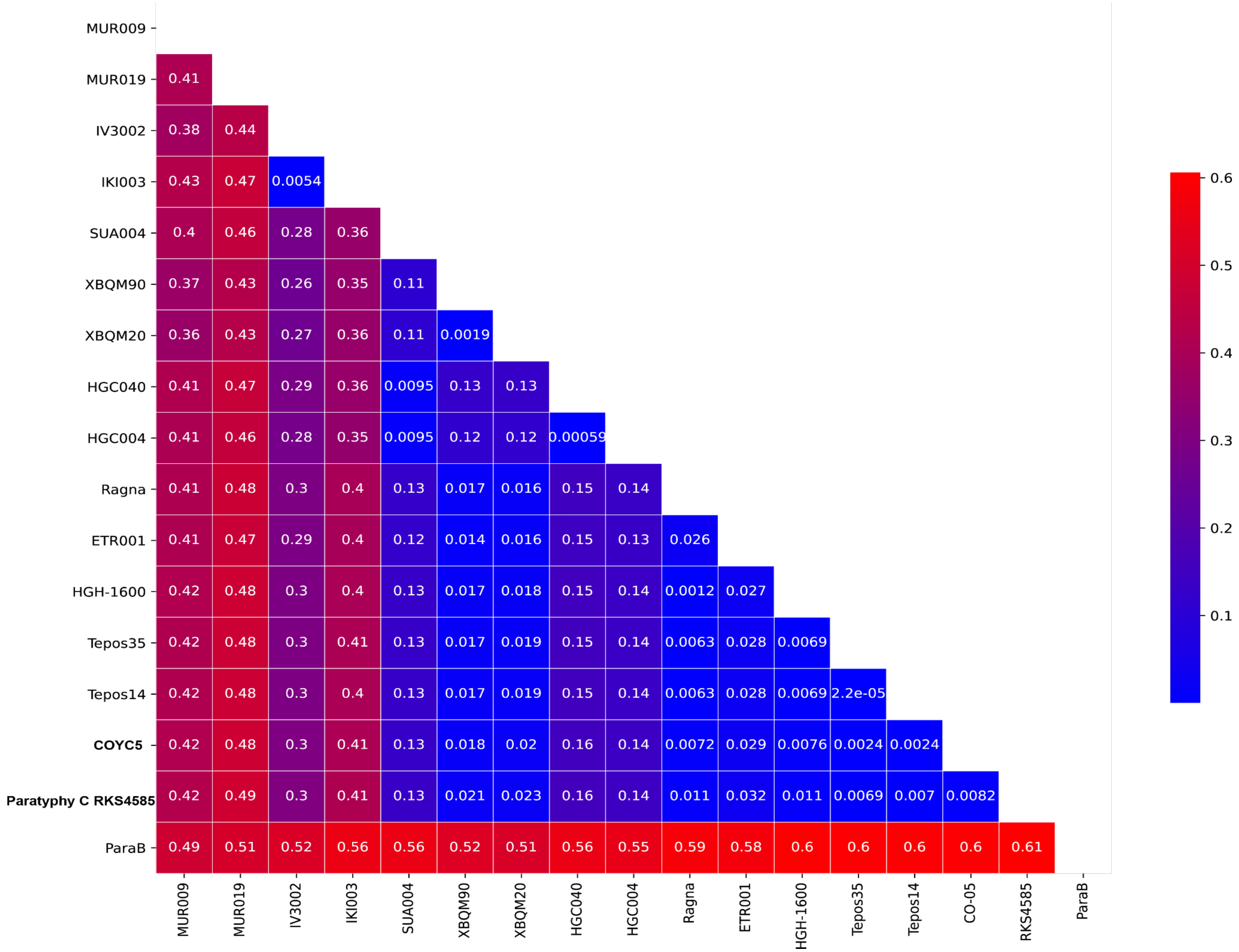


Figure S6. Pairwise distances between COYC5 and ancient *Salmonella enterica* Paratyphi C genomes are visualized in a symmetric matrix and plotted as a heatmap.

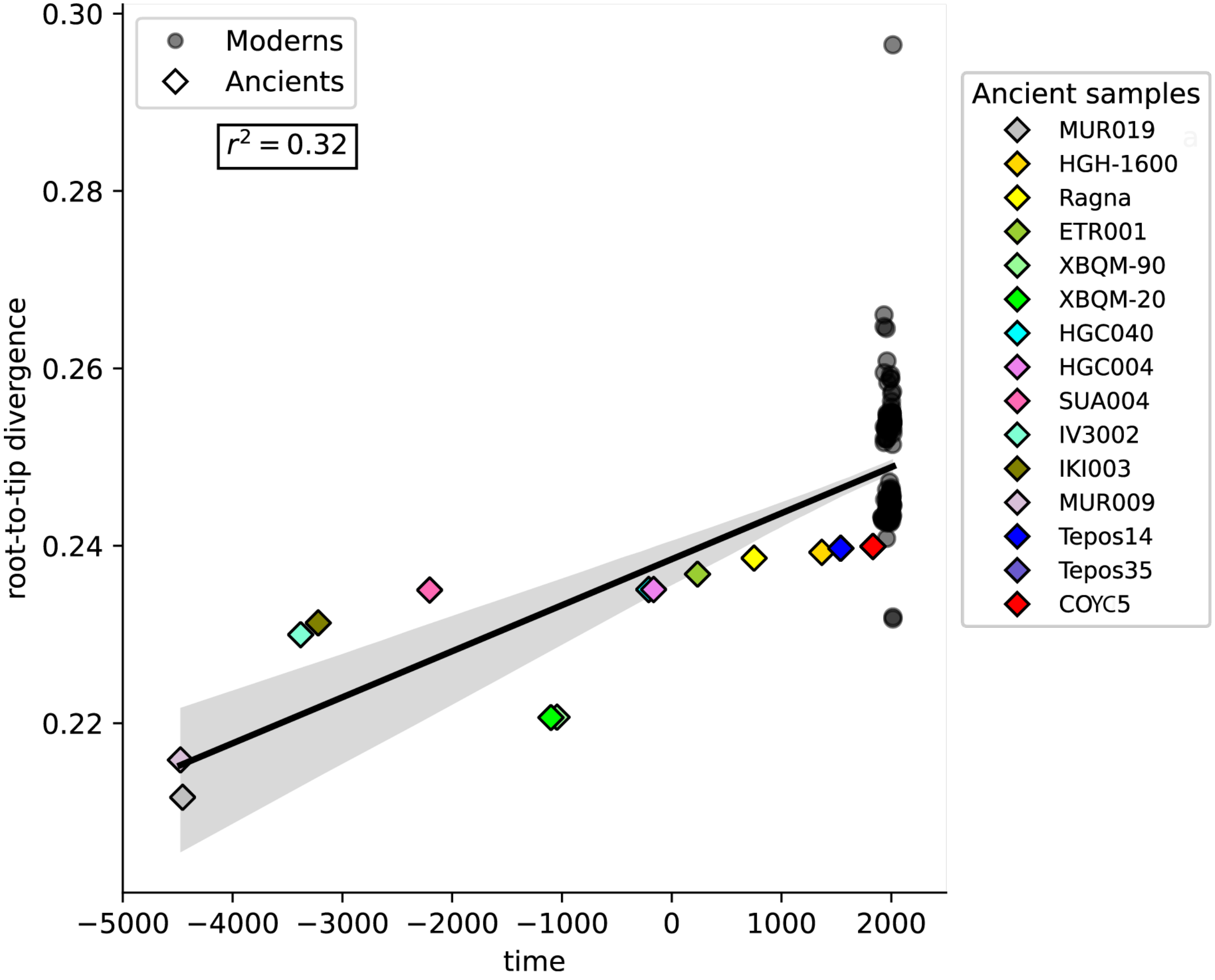


Figure S7. Assessment of the temporal signal using *TempEst*. Root-to-tip genetic distances were plotted against sampling dates.

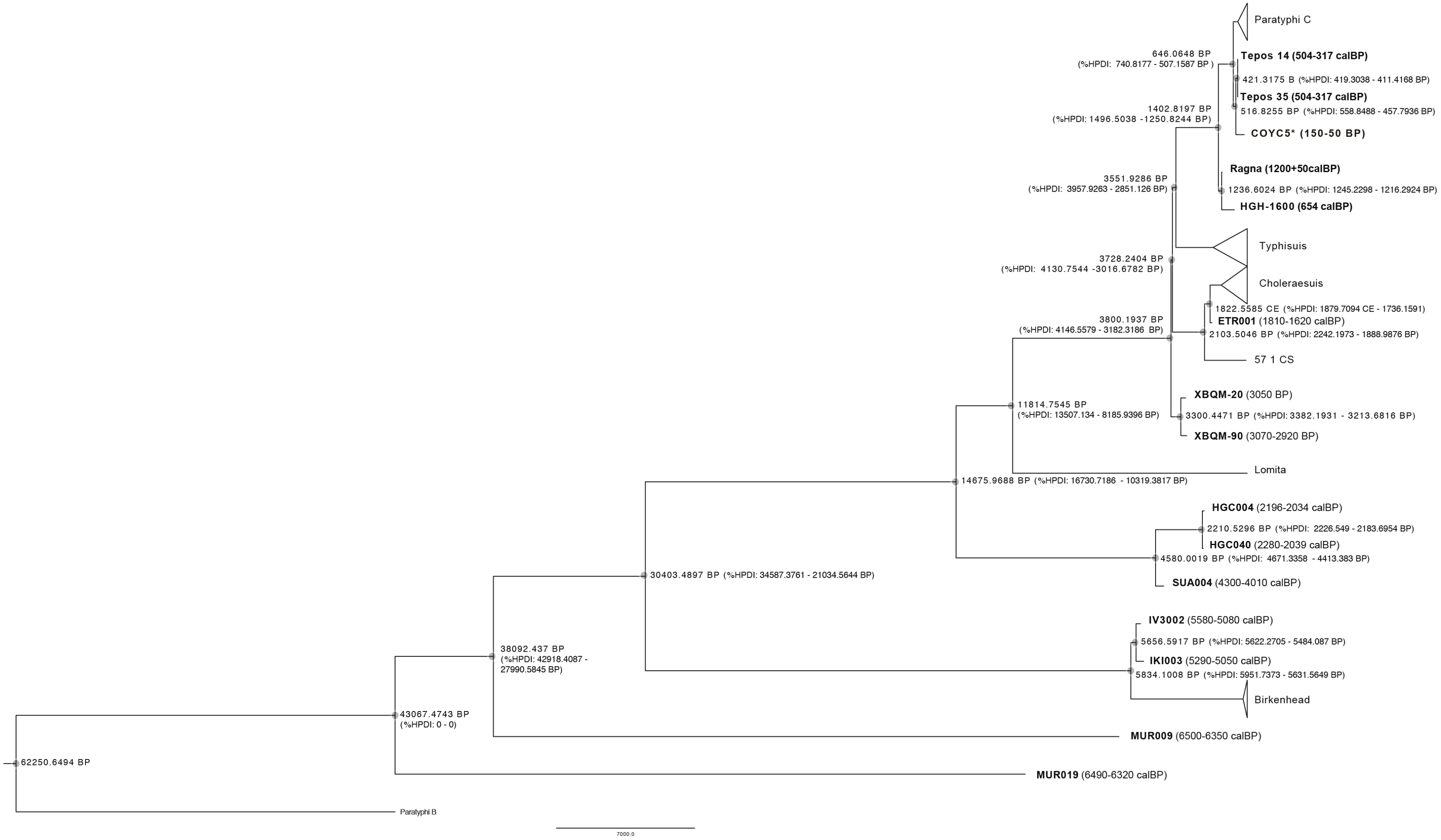


Figure S8. Bayesian time-calibrated phylogenetic tree including COYC5 individual, 14 ancient *Salmonella enterica* genomes, and 221 modern genomes analyzed in this study. Node labels indicate posterior probabilities and 95% highest posterior density intervals (%HPDI) for estimated divergence times at major nodes, expressed in BP years.

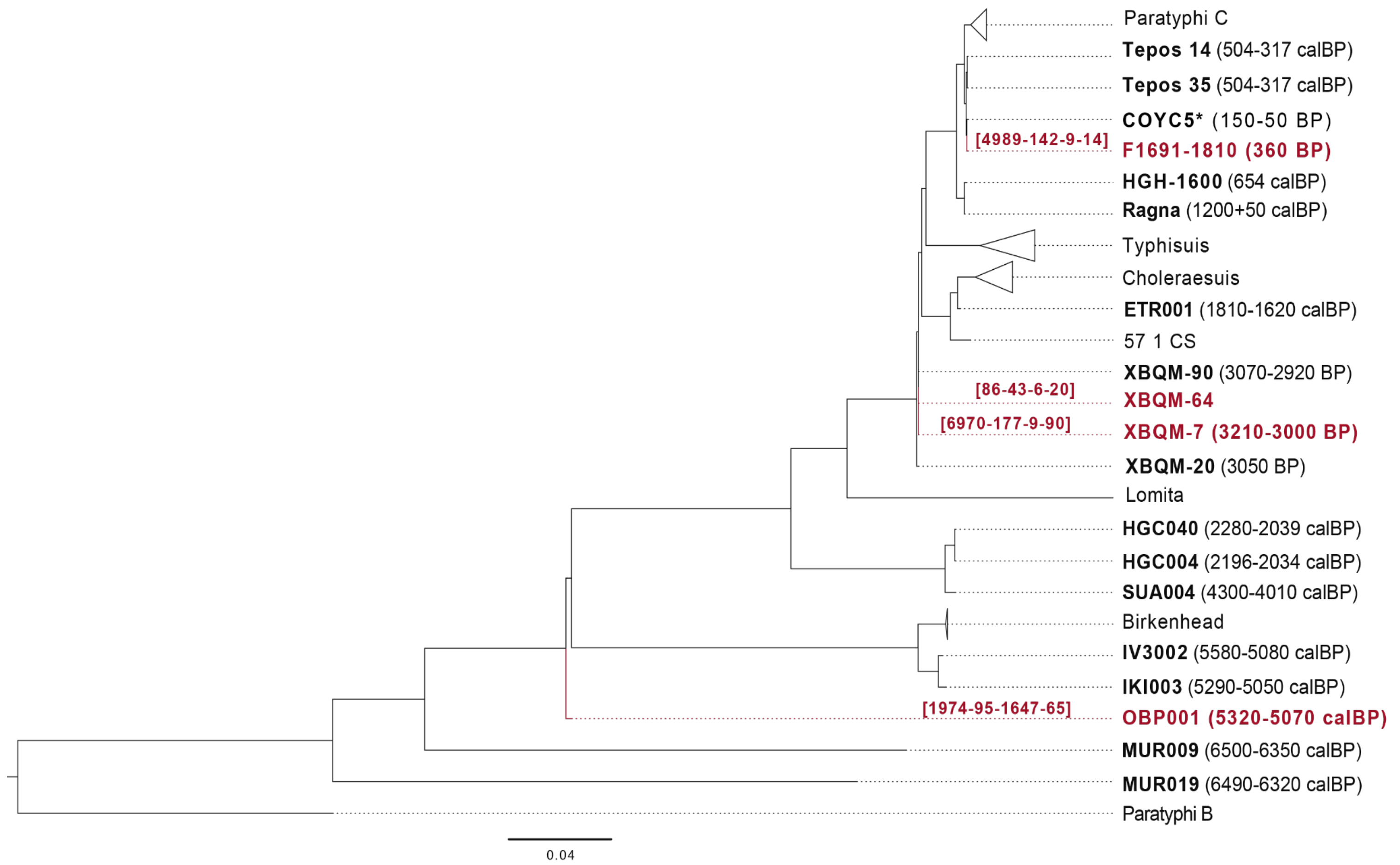


Figure S9. Phylogenetic placement using *PathPhynder*, considering four low coverage ancient samples previously reported: F1691-180 (0.3X) (de-Dios et al. 2021); XBQM-64 (0.19-0.92 X), XBQM-7 (1.2X) (Wu et al. 2021); and OBP001 (1.2X) (Key et al. 2020). Each label consists of four values separated by hyphens, corresponding to the number of supporting markers above the branch – conflicting markers above the branch – supporting markers on the branch – conflicting markers on the branch. A placement is considered reliable when the values in the first and third positions are higher than those in the second and fourth. Conversely, if the second and fourth values exceed the first and third, the placement is not reliable.

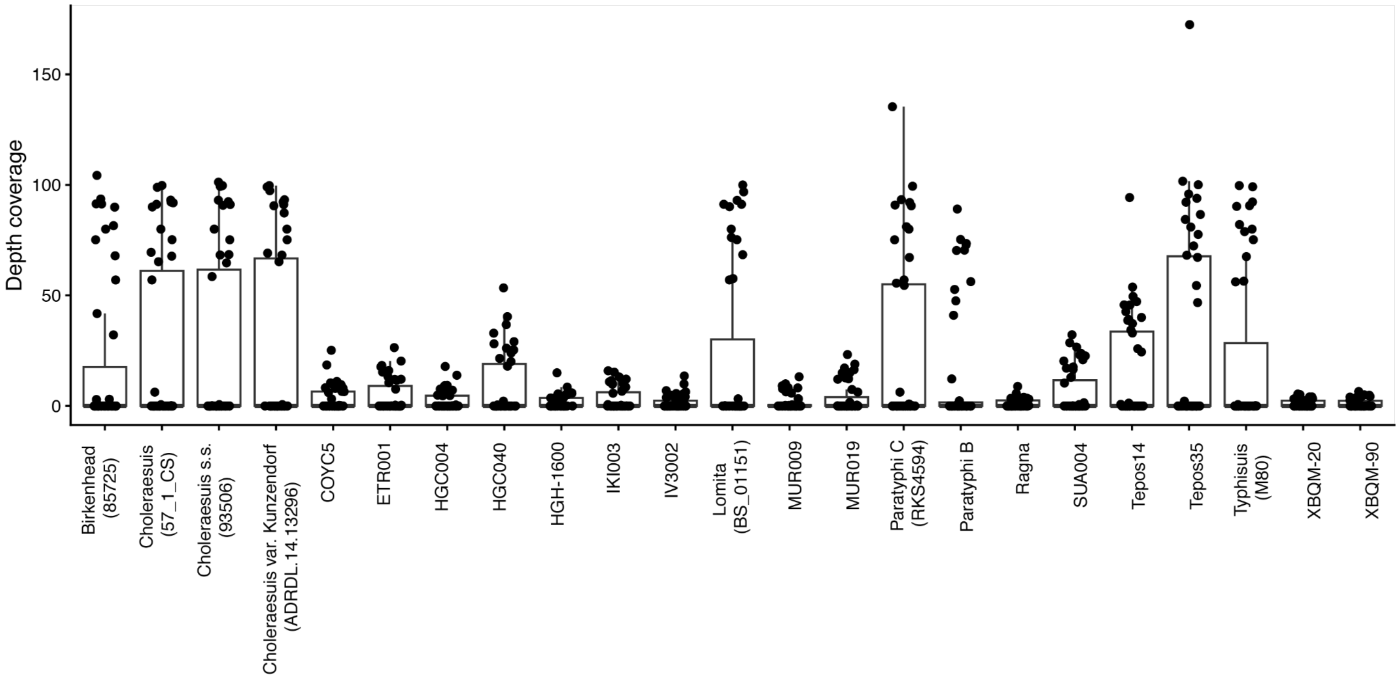


Figure S10. Boxplot of the average raw depth values of the *Salmonella* pathogenicity islands (SPI-1 to SPI-14), including virulence genes in SPI-6 and SPI-7.

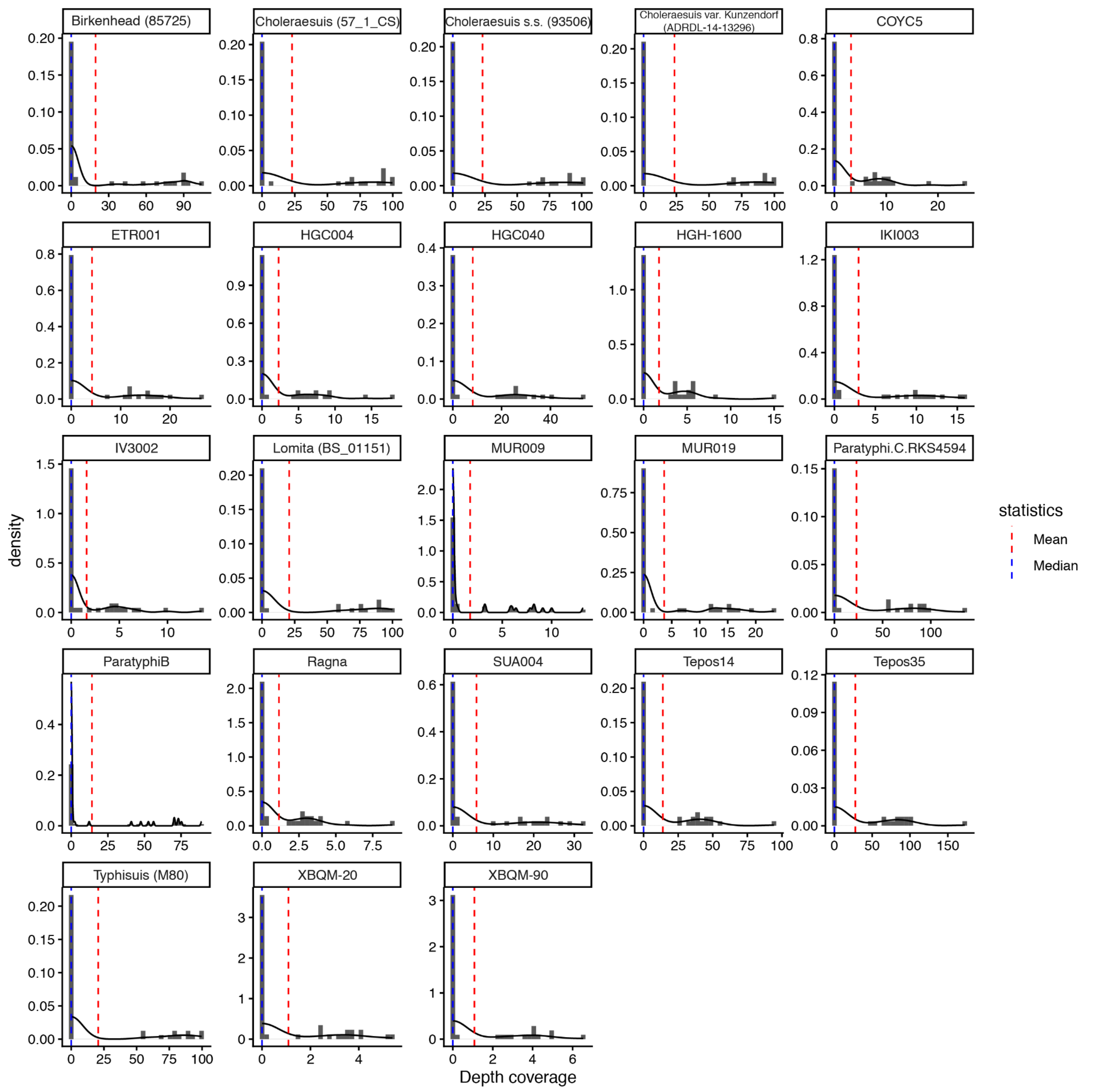


Figure S11. Histograms showing the distribution of the raw average depth values of the *Salmonella* pathogenicity islands (SPI-1 to SPI-14), including virulence genes in SPI-6 and SPI-7. The mean and median for each sample are indicated with vertical lines.

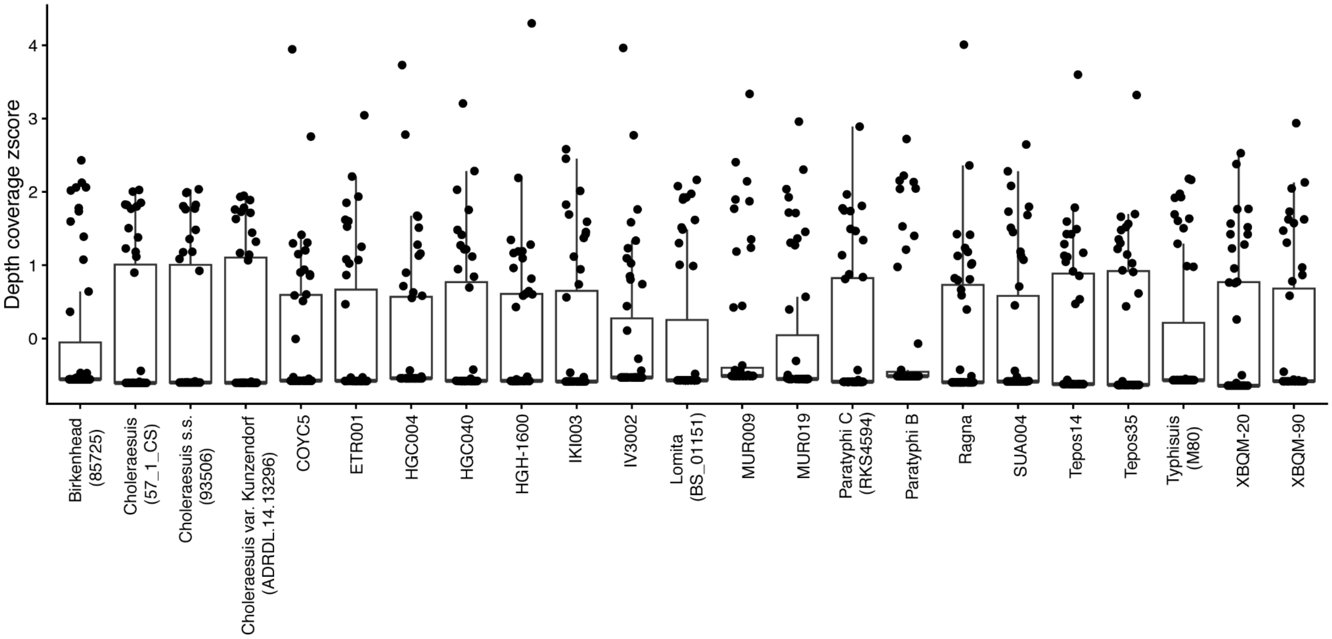


Figure S12. Boxplot of the normalized average depth values of the *Salmonella* pathogenicity islands (SPI-1 to SPI-14), including virulence genes in SPI-6 and SPI-7.

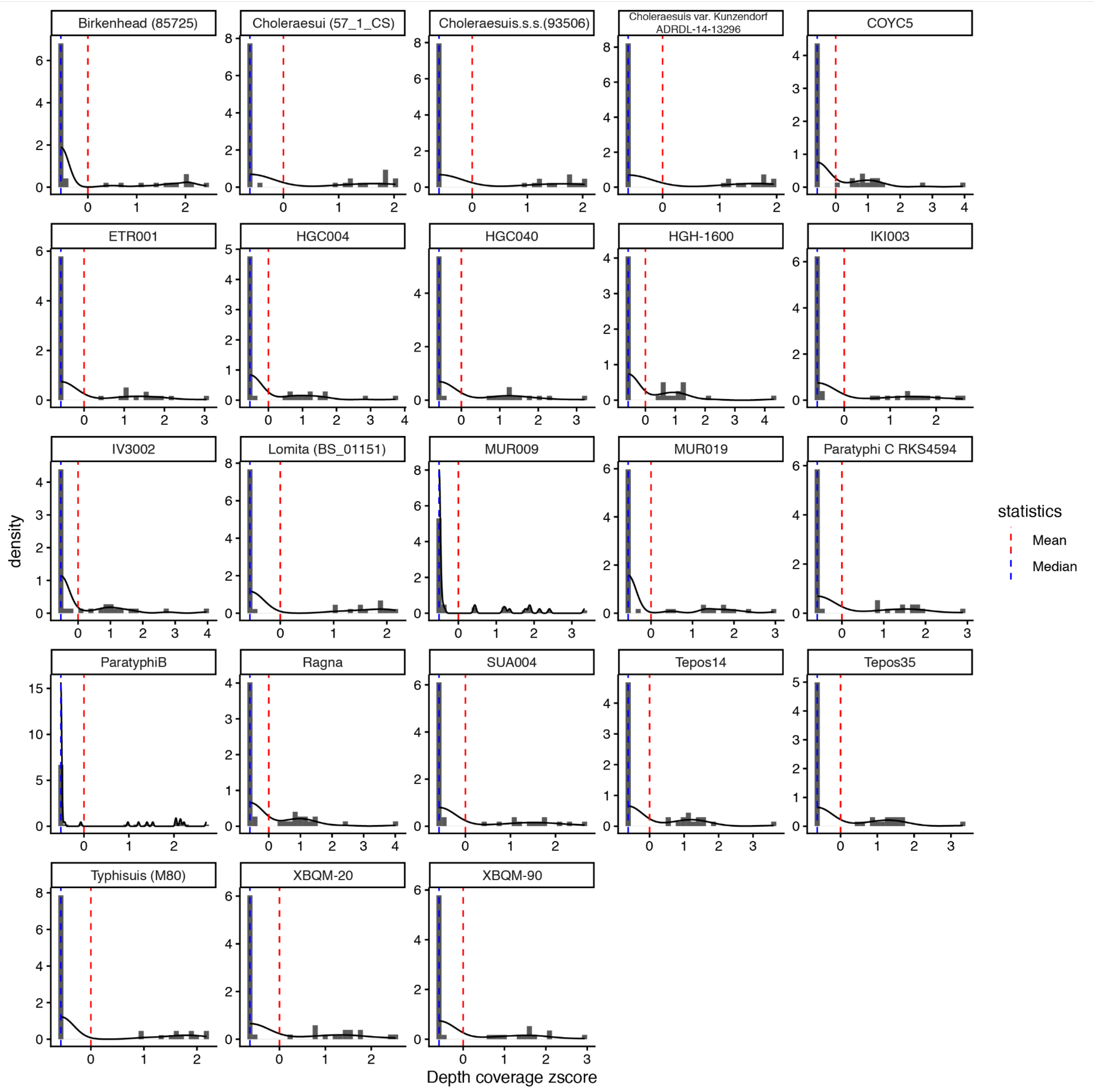


Figure S13. Histograms showing the distribution of normalized average depth values for the *Salmonella* pathogenicity islands (SPI-1 to SPI-14), including virulence genes in SPI-6 and SPI-7.The mean and median for each sample are indicated with vertical lines.

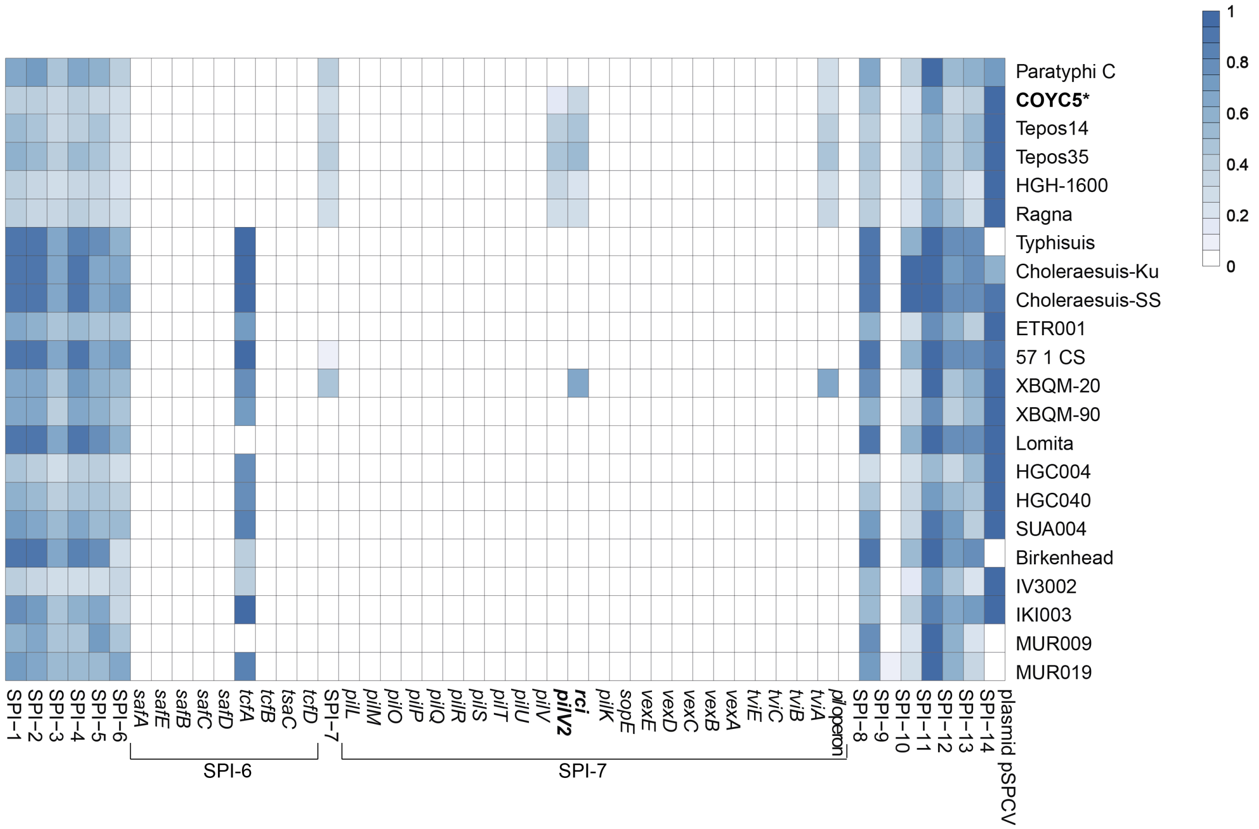


Figure S14. Heatmap showing the presence and absence of *Salmonella* pathogenicity islands (SPI-1 to SPI-14), including key virulence genes in SPI-6 and SPI-7, based on the PAIDB database (Yoon et al., 2015). The genes related to the shufflon mechanism are highlighted in bold within the SPI-7. Values were scaled between 0 and 1 using min–max normalization to enable comparison across individuals.

### References

AlQahtani, S. J., Hector, M. P., & Liversidge, H. M. (2010). Brief communication: The London atlas of human tooth development and eruption. American Journal of Physical Anthropology, 142(3), 481–490. https://doi.org/10.1002/ajpa.21258

de Dios, T., Carrión, P., Olalde, I., Llovera Nadal, L., Lizano, E., Pàmies, D., Marques-Bonet, T., Balloux, F., van Dorp, L., & Lalueza-Fox, C. (2021). Salmonella enterica from a soldier from the 1652 Siege of Barcelona (Spain) supports historical transatlantic epidemic contacts. iScience, 24(9), 103021. https://doi.org/10.1016/j.isci.2021.103021

Goodman, A. H., & Martin, D. L. (2002). Reconstructing health profiles from skeletal remains. In J. C. Rose & R. H. Steckel (Eds.), The backbone of history: Health and nutrition in the Western Hemisphere (pp. 11–60). Cambridge University Press.

Gu, Z., Gu, L., Eils, R., Schlesner, M., & Brors, B. (2014). Circlize implements and enhances circular visualization in R. Bioinformatics, 30(19), 2811–2812. https://doi.org/10.1093/bioinformatics/btu393

Key, F. M., Posth, C., Esquivel-Gomez, L. R., Hübler, R., Spyrou, M. A., Neumann, G. U., Furtwängler, A., et al. (2020). Emergence of human-adapted Salmonella enterica is linked to the Neolithization process. Nature Ecology & Evolution, 4(3), 324–333. https://doi.org/10.1038/s41559-020-1111-0

Klales, A. R. (2020). MorphoPASSE: Morphological pelvis and skull sex estimation program. In Sex estimation of the human skeleton (pp. 271–278). Elsevier.

Knüsel, C. J., Göggel, S., & Lucy, D. (1997). Comparative degenerative joint disease of the vertebral column in the medieval monastic cemetery of the Gilbertine Priory of St. Andrew, Fishergate, York, England. American Journal of Physical Anthropology, 103(4), 481–495. https://doi.org/10.1002/(SICI)1096-8644(199708)103:4

Larsen, C. S. (2018). Bioarchaeology in perspective: From classifications of the dead to conditions of the living. American Journal of Physical Anthropology, 165(4), 865–878. https://doi.org/10.1002/ajpa.23322

Lottering, N., MacGregor, D. M., Alston, C. L., & Gregory, L. S. (2015). Ontogeny of the spheno-occipital synchondrosis in a modern Queensland, Australian population using computed tomography. American Journal of Physical Anthropology, 157(1), 42–57. https://doi.org/10.1002/ajpa.22687

Ortner, D. J. (2003). Identification of pathological conditions in human skeletal remains (2nd ed.). Academic Press.

Villotte, S., & Knüsel, C. J. (2013). Understanding entheseal changes: Definition and life course changes. International Journal of Osteoarchaeology, 23(2), 135–146. https://doi.org/10.1002/oa.2289

White, T., & Folkens, P. (2005). The human bone manual. Academic Press.

Wu, X., Ning, C., Key, F. M., Andrades Valtueña, A., Lankapalli, A. K., Gao, S., Yang, X., et al.

(2021). A 3,000-year-old, basal S. enterica lineage from Bronze Age Xinjiang suggests spread along the Proto-Silk Road. PLoS Pathogens, 17(9), e1009886. https://doi.org/10.1371/journal.ppat.1009886

Yoon, S. H., Park, Y.-K., & Kim, J. F. (2015). PAIDB v2.0: exploration and analysis of pathogenicity and resistance islands. Nucleic Acids Research, 43(Database issue), D624–D630.
